# Virological characteristics of the SARS-CoV-2 XBB variant derived from recombination of two Omicron subvariants

**DOI:** 10.1101/2022.12.27.521986

**Authors:** Tomokazu Tamura, Jumpei Ito, Keiya Uriu, Jiri Zahradnik, Izumi Kida, Hesham Nasser, Maya Shofa, Yoshitaka Oda, Spyros Lytras, Naganori Nao, Yukari Itakura, Sayaka Deguchi, Rigel Suzuki, Lei Wang, MST Monira Begum, Masumi Tsuda, Yusuke Kosugi, Shigeru Fujita, Kumiko Yoshimatsu, Saori Suzuki, Hiroyuki Asakura, Mami Nagashima, Kenji Sadamasu, Kazuhisa Yoshimura, Yuki Yamamoto, Tetsuharu Nagamoto, Gideon Schreiber, The Genotype to Phenotype Japan (G2P-Japan) Consortium, Terumasa Ikeda, Takasuke Fukuhara, Akatsuki Saito, Shinya Tanaka, Keita Matsuno, Kazuo Takayama, Kei Sato

## Abstract

In late 2022, the SARS-CoV-2 Omicron subvariants have highly diversified, and XBB is spreading rapidly around the world. Our phylogenetic analyses suggested that XBB emerged by recombination of two co-circulating BA.2 lineages, BJ.1 and BM.1.1.1 (a progeny of BA.2.75), during the summer of 2022 around India. *In vitro* experiments revealed that XBB is the most profoundly resistant variant to BA.2/5 breakthrough infection sera ever and is more fusogenic than BA.2.75. Notably, the recombination breakpoint is located in the receptor-binding domain of spike, and each region of recombined spike conferred immune evasion and augmented fusogenicity to the XBB spike. Finally, the intrinsic pathogenicity of XBB in hamsters is comparable to or even lower than that of BA.2.75. Our multiscale investigation provided evidence suggesting that XBB is the first documented SARS-CoV-2 variant increasing its fitness through recombination rather than single mutations.

## Introduction

The SARS-CoV-2 Omicron variant is the current variant of concern since the end of 2021 (ref.^1^). As of December 2022, recently emerging Omicron subvariants are under convergent evolution: recently emerging variants acquired substitutions at the same residues of the spike (S) protein, such as R346, K444, L452, N460, and F486^2, 3^. For instance, Omicron BQ.1.1 variant, which is a descendant of Omicron BA.5 and is currently becoming predominant in the Western countries^1^, possesses all convergent substitutions, such as R346T, K444T, L452R, N460K, and F486V. Recent studies including ours suggested that L452R^4–9^, N460K^2, 6, 10, 11^, and R346T^2^ increase the binding affinity of SARS-CoV-2 S protein to human angiotensin-converting enzyme 2 (ACE2), the receptor for viral infection, while R346T^12, 13^, K444T^12^ and F486V^2, 4, 5, 12, 14, 15^ contribute to evade antiviral humoral immunity induced by vaccination and natural SARS-CoV-2 infection. Similar to the observations in BA.5 (ref.^5^) and BA.2.75 (ref.^10^), combinational substitutions in S protein (1) to evade antiviral humoral immunity in exchange for the decrease of ACE2 binding affinity (e.g., F486V) and (2) to enhance ACE2 binding affinity to compensate the decreased affinity by immune evasion substitution (e.g., L452R and N460K) has been frequently observed in recently emerging Omicron subvariants including BQ.1.1. These observations suggest that acquiring these two types of substitutions in the S protein is a trend for recently emerging Omicron subvariants to spread more efficiently than prior ones.

In addition to the diversification and subsequent convergent evolution of emerging Omicron subvariants (e.g., BQ.1.1), a recombinant variant, called XBB, has recently emerged. The Omicron XBB variant likely originated by the recombination of two BA.2 descendants, BJ.1 and BM.1.1.1 (a progeny of BA.2.75)^16^. While the BQ.1 lineage is becoming predominant in Europe, XBB has become predominant in India and Singapore and is spreading several countries^17^. As of October 28, 2022, the WHO classifies XBB as an Omicron subvariant under monitoring^1^. Recent studies including ours have revealed the virological features of BQ.1 (refs.^2, 13, 18^). However, the features of XBB, another Omicron subvariants of concern, are not fully elucidated. In this study, we elucidated the virological characteristics of XBB, particularly its transmissibility, immune resistance, ACE2 binding affinity, infectivity, fusogenicity and intrinsic pathogenicity in a hamster model.

## Results

### Evolution and epidemics of the XBB variant

As of December 2022, most of the prevalent Omicron lineages, including BA.5, are descendants of BA.2 (**Fig. 1a**). Of these, highly diversified BA.2 subvariants, such as BA.2.75 and BJ.1, have emerged in South Asia and are referred to as the second-generation BA.2 variants (**Fig. 1a**). Recently, the XBB variant emerged as a recombinant lineage between the second generation BA.2 variants, BJ.1 and BM.1.1.1 (BA.2.75.3.1.1.1; a descendant of BA.2.75)^16^ (**Fig. 1a**). XBB harbors the S substitutions R346T, N460K, and F486S, which have been convergently acquired during the Omicron evolution (**Fig. 1b**)^2^. To trace the recombination event that led to the emergence of the XBB variant, we retrieved all SARS-CoV-2 sequences deposited to GISAID (as of October 3, 2022) with PANGO lineage designation matching BJ.1, BM.1, XBB, and all their descendant lineages (including BM.1.1, BM.1.1.1, and XBB.1). Recombination analysis on the aligned set of sequences, using RDP5 (ref.^19^), robustly picks up a single recombination breakpoint unique to all XBB sequences at the genomic position 22,920 (matching the Wuhan-Hu-1 reference genome) (**Fig. 1c**). No evidence of recombination was found in the BJ.1 and BM.1 sequences in the dataset. Consistent with the result of the RDP5 analysis, the visual inspection of the nucleotide differences between the consensus sequences of XBB, BJ.1, and BM.1 (including BM.1.1 and BM.1.1.1) clearly illustrates that the XBB identity to BJ.1 ends at the genome position 22,942 and the XBB identity to BM.1 starts after position 22,896 (**Fig. 1c**). Together, our recombination analysis suggests that the recombination breakpoint is between positions 22,897 and 22,941, within the receptor binding domain (RBD) of S protein (corresponding to amino acid positions 445-460) (**Fig. 1c**).

**Fig. 1.**
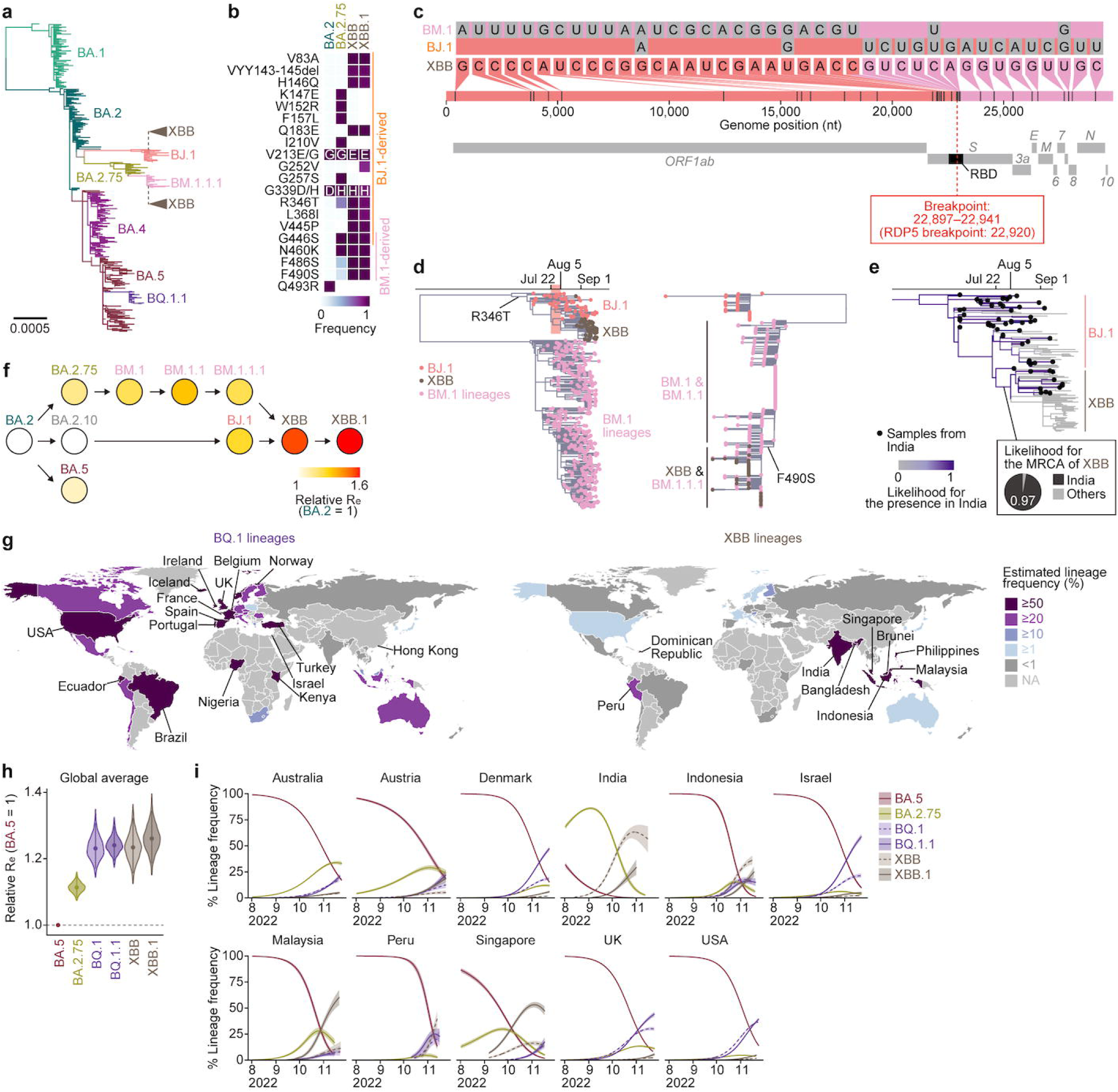
Phylogenetic and epidemic analyses of the XBB lineage. **a,** Phylogenetic tree of representative sequences from PANGO lineages of interest: BA.1, BA.2, BA.4, BA.5, BA.2.75, BJ.1, and BM.1.1.1, rooted on a B.1.1 outgroup (not shown). The recombinant parents of XBB are annotated on the tree as cartoon clades. **b**, Amino acid differences in the S proteins of Omicron lineages. **c,** Nucleotide differences between the consensus sequences of the BJ.1, BM.1 (including BM.1.1/BM.1.1.1) lineages and the XBB (including XBB.1) lineage, visualized with snipit (https://github.com/aineniamh/snipit). **d,** Maximum likelihood time-calibrated phylogeny of the 5’ non-recombinant segment (1–22,920) of the XBB variant (left) and non-calibrated phylogeny of the 3’ non-recombinant segment (22,920–29,903) (right). Both trees are rooted on a BA.2 outgroup (not shown). **e,** Ancestral state reconstruction of the circulated regions of viruses. The time-calibrated tree shown in **Fig. 1d, left** is inherited in this analysis. Phylogeny related to XBB is only shown. Black circles on tips indicate samples from India. Branch color indicates the ancestral likelihood of the presence in India. In addition, the ancestral likelihood value for the MRCA of XBB lineages is shown as a pie chart. See also **Extended Data Fig. S1a**. **f,** Relative R_e_ values for viral lineages in India, assuming a fixed generation time of 2.1 days. The R_e_ of BA.2 is set at 1. Dot color indicates the posterior mean of the R_e_, and an arrow indicate phylogenetic relationship. See also **Extended Data Fig. S1b**. **g,** Difference in the circulated regions between BQ.1 and XBB lineages. Estimated lineage frequency as of November 15^th^, 2022 in each country is 50% and ≥20% frequencies are annotated for the BQ.1 and XBB lineages, respectively. **h,** Relative R_e_ values for viral lineages, assuming a fixed generation time of 2.1 days. The R_e_ value of BA.5 is set at 1. The posterior (violin), posterior mean (dot), and 95% Bayesian confidential interval (CI; line) are shown. The global average values estimated by a hierarchical Bayesian model^23^ are shown. See also **Extended Data Fig. S1c**. **i,** Estimated lineage dynamics in each country where BQ.1 and XBB lineages cocirculated.

We then split the sequence alignment at position 22,920 to determine the evolutionary history of each non-recombinant segment of the XBB genomes. The phylogenetic reconstructions recapitulate the recombination results, with the 5’ end major parental sequence being derived from the BJ.1 clade and the 3’ end minor parental sequence from the BM.1.1.1 clade (**Fig. 1d**). Using the longer 5’ end non-recombinant part of these genomes, we estimated the emergence date of XBB based on the inferred root-to-tip regression (see **Methods**) (**Fig. 1d**). This analysis suggests that the XBB clade stemmed from the major BJ.1 clade in the span from July 22 to August 5, 2022 (**Fig. 1d**). Furthermore, we inferred the emergence region of XBB by a discrete ancestral state reconstruction utilizing a Markov model (see **Methods**). Most of the earlier isolates of XBB were sampled in India, and our analysis suggests that the ancestral state (i.e., the circulating region) of the most recent common ancestor (MRCA) of XBB sequences is likely India (**Fig. 1e and Extended Data Fig. S1b**). Together, our analyses suggest that XBB emerged through the recombination of two co-circulating lineages, BJ.1 and BM.1.1.1, during the summer of 2022 in India or neighboring countries.

To trace the shift of viral fitness during the evolution of Omicron leading to the emergence of XBB, we estimated the effective reproduction number (R_e_) of XBB-related variants based on the epidemic data of SARS-CoV-2 in India (from June 1 to November 15, 2022) (**Fig. 1f and Extended Data Fig. S1c and Supplementary Table 1**). BJ.1 and BM.1/BM.1.1/BA.1.1.1 showed higher R_e_ compared with their parental lineages, BA.2.10 and BA.2.75, respectively. Furthermore, the R_e_ value of XBB is 1.23- and 1.20-times higher than those of the parental BJ.1 and BM.1.1.1, respectively (**Fig. 1f and Extended Data Fig. S1c and Supplementary Table 1**). Importantly, this is the first documented example of a SARS-CoV-2 variant increasing its fitness through recombination rather than single mutations.

As of December 2022, two viral lineages are expanding their epidemics around the world: BQ.1 lineages and XBB lineages. To investigate the prevalence of these two lineages in various geographic regions, we estimated the epidemic frequency of each variant as of November 15, 2022, in each county (**Fig. 1g and Supplementary Table 2**). BQ.1 lineages have spread and reached dominance in European, American, and African countries, probably reflecting the likelihood that BQ.1 emerged from the African continent^20^ (**Fig. 1g**). On the other hand, XBB lineages have spread and reached dominance in South and Southeast Asian countries, such as India, Singapore, and Indonesia, reflecting the fact that XBB originated around India (**Fig. 1e,g**). Furthermore, we constructed a hierarchical Bayesian model and estimated the global average and country-specific R_e_ values of XBB lineages according to the epidemic data of countries where XBB lineages cocirculated with BQ.1 lineages (**Fig. 1h,i, Extended Data Fig. S1d and Supplementary Table 3**). Our analysis shows that the R_e_ values of XBB and XBB.1 (i.e., XBB harboring S:G252V) are 1.24 and 1.26-times higher than that of BA.5 and are comparable with those of BQ.1 and BQ.1.1 (**Fig. 1h and Extended Data Fig. S1d**). Together, our analyses show that both BQ.1 and XBB lineages, exhibiting a similar advantage in viral fitness, are becoming predominant in the Western and Eastern Hemispheres, respectively.

### Immune resistance of XBB

To investigate the virological features of XBB, we first evaluated the immune resistance of XBB using HIV-1-based pseudoviruses. In the present study, we used the major S haplotype of XBB lineages as of October 3, 2022, corresponding to the S protein of XBB.1, for the following experiments (hereafter we simply refer to XBB.1 as XBB). In the case of breakthrough BA.2 infection sera, BA.2.75 did not exhibit significant resistance when compared to BA.2 (**Fig. 2a**), which is consistent with our prior study^10^. In contrast, we found that XBB exhibits a profound (30-fold) resistance to breakthrough BA.2 infection sera (*P*=0.0002, **Fig. 2a**). To determine amino acid substitutions conferring the profound resistance to the breakthrough anti sera, we constructed the BA.2 S mutants which harbor respective single substitutions present in XBB. We did not analyze the substitutions present also in BA.2.75 (e.g., G446S) since we have already analyzed these substitutions in our previous study^10^. As shown in **Fig. 2a**, several substitutions such as V83A (2.1-fold, *P*=0.0034), Y144del (2.9-fold, *P*=0.0002), Q183E (2.0-fold, *P*=0.0039), R346T (2.1-fold, *P*=0.0005), L368I (1.8-fold, *P*=0.042), V445P (2.1-fold, *P*=0.0002), F486S (3.0-fold, *P*=0.0002), and F490S (2.7-fold, *P*=0.024) conferred significant resistance to breakthrough BA.2 infection sera. Because the immune resistance conferred by respective substitution is relatively minor when compared to the resistance of XBB (**Fig. 2a**), our data suggest that multiple substitutions in the XBB S cooperatively contribute to the resistance against humoral immunity induced by breakthrough BA.2 infection.

**Fig. 2.**
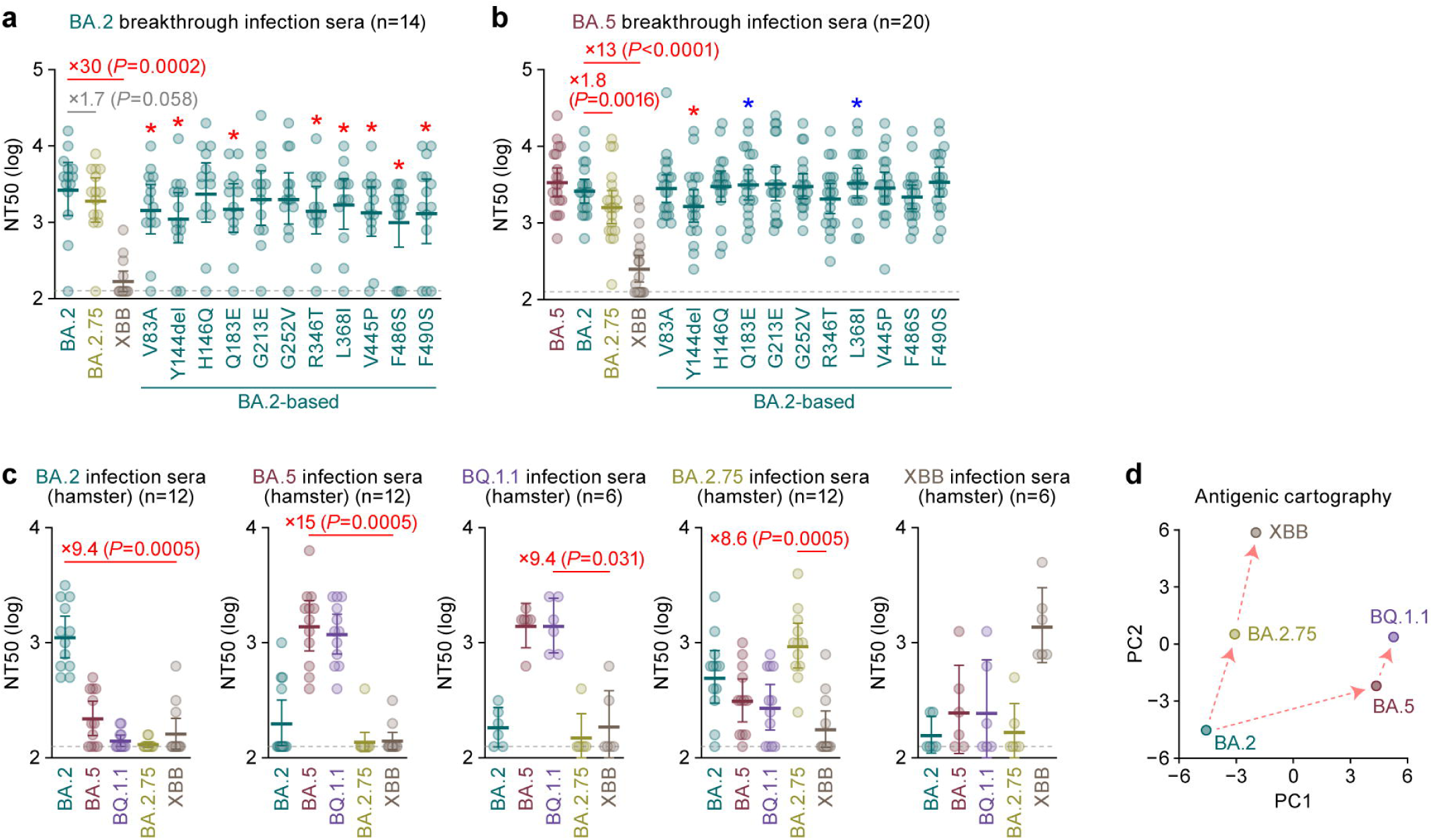
Immune resistance of XBB. Neutralization assays were performed with pseudoviruses harboring the S proteins of BA.2, BA.5, BQ.1.1, BA.2.75 and XBB. The BA.2 S-based derivatives are included in **a and b**. The following sera were used. **a,b,** Convalescent sera from fully vaccinated individuals who had been infected with BA.2 after full vaccination (9 2-dose vaccinated and 5 3-dose vaccinated. 14 donors in total) (**a**), and BA.5 after full vaccination (2 2-dose vaccinated donors, 17 3-dose vaccinated donors and 1 4-dose vaccinated donors. 20 donors in total) (**b**). **c,** Sera from hamsters infected with BA.2 (12 hamsters), BA.5 (12 hamsters), BQ.1.1 (6 hamsters), BA.2.75 (12 hamsters), and XBB (6 hamsters). **d,** Antigenic cartography based on the results of neutralization assays using hamster sera (**Fig. 2c**). Assays for each serum sample were performed in triplicate to determine the 50% neutralization titer (NT_50_). Each dot represents one NT_50_ value, and the geometric mean and 95% CI are shown. Statistically significant differences were determined by two-sided Wilcoxon signed-rank tests. The *P* values versus BA.2 (**a**), BA.5 (**b**), or XBB (**c**) are indicated in the panels. For the BA.2 derivatives (**a and b**), statistically significant differences (*P* < 0.05) versus BA.2 are indicated with asterisks. Red and blue asterisks, respectively, indicate decreased and increased NT50s. The horizontal dashed line indicates the detection limit (120-fold). Information on the convalescent donors is summarized in **Supplementary Table 4**.

Consistent with our previous study^10^, BA.2.75 showed a statistically significant (1.8-fold) resistance to breakthrough BA.5 infection sera when compared to BA.2 (*P*=0.0016, **Fig. 2b**). Moreover, XBB exhibited a profound (13-fold) resistance to breakthrough BA.5 infection sera (*P*<0.0001, **Fig. 2b**). Neutralization assay using the pseudoviruses with BA.2 derivatives revealed that the Y144del mutation (1.8-fold, *P*=0.016) resulted in the resistance to breakthrough BA.5 infection sera (**Fig. 2b**). Furthermore, in our previous study, we showed that G446S, which is a common substitution of BA.2.75 and XBB, conferred immune resistance to breakthrough BA.5 infection sera^10^. Together, these observations suggest that these two mutations (Y144del and G446S) cooperatively contribute to the resistance against humoral immunity induced by breakthrough BA.5 infection.

To further evaluate the antigenicity of XBB S, we used the sera obtained from infected hamsters at 16 days post infection (d.p.i.). As shown in **Fig. 2c**, XBB exhibited profound resistance to the sera obtained from hamsters infected with BA.2, BA.5, BQ.1.1, and BA.2.75. Moreover, XBB infection hamster sera exhibited remarkable antiviral effect against only XBB (**Fig. 2c**). The cartography based on the neutralization dataset using hamster sera (**Fig. 2c**) showed that the cross-reactivity of each Omicron subvariant is correlated to their phylogenetic relationship (**Fig. 1a**). and the antigenicity of XBB is distinct from the other Omicron subvariants tested (**Fig. 2d**). These observations suggest that XBB is antigenically different from the other Omicron subvariants including BQ.1.1, and therefore, remarkably evades the BA.2/5 infection-induced herd immunity in the human population.

### ACE2 binding affinity of XBB S

We then evaluated the features of XBB S that potentially affect viral infection and replication. Yeast surface display assay^2, 10, 21, 22^ showed that the binding affinity of XBB S RBD to human ACE2 receptor (1.00±0.069) is significantly lower than that of ancestral BA.2 S RBD (1.49±0.054) (**Fig. 3a**). As described above (**Fig. 1a–c**), the four RBD substitutions in XBB compared to BA.2, D339H, G446S, N460K and R493Q, are common to BA.2.75 since a part of RBD of XBB S is derived from the BA.2.75/BM.1 lineage. In our previous studies^2, 10^, we demonstrated that the N460K substitution augments ACE2 binding affinity. To address whether other substitutions in the XBB S affect the binding affinity of S RBD to human ACE2, we prepared a repertoire of BA.2 S RBD that possesses an XBB-specific substitution compared to BA.2. Consistent with our recent study^2, 10^, the R346T substitution, which is common in both XBB and BQ.1.1, significantly increased the binding affinity of BA.2 S RBD to human ACE2 (**Fig. 3a**). Additionally, the L368I substitution also augmented ACE2 binding affinity (**Fig. 3a**). On the other hand, the F486S substitution significantly decreased ACE2 binding affinity (**Fig. 3a**). Because the F486V substitution also decreased ACE2 binding affinity^5^, our data suggest that the amino acid substitution at F486 leads to attenuated ACE2 binding affinity. Our results suggested that the enhanced binding affinity of XBB S RBD compared to BA.2 S RBD is attributed to at least three substitutions in the RBD: R346T, L368I and N460K^2, 10^. Nevertheless, the K_D_ value of XBB S RBD was clearly higher than that of BA.2.75 S RBD (0.18±0.069) (**Fig. 3a**). In our prior study^10^, we showed that the D339H substitution contributes to the augment of ACE2 binding affinity only when the backbone is the BA.2.75 S RBD. Therefore, the profound binding affinity of BA.2.75 S RBD to human ACE2 would be attributed to the conformation that is composed of multiple substitutions in the BA.2.75 S RBD.

**Fig. 3.**
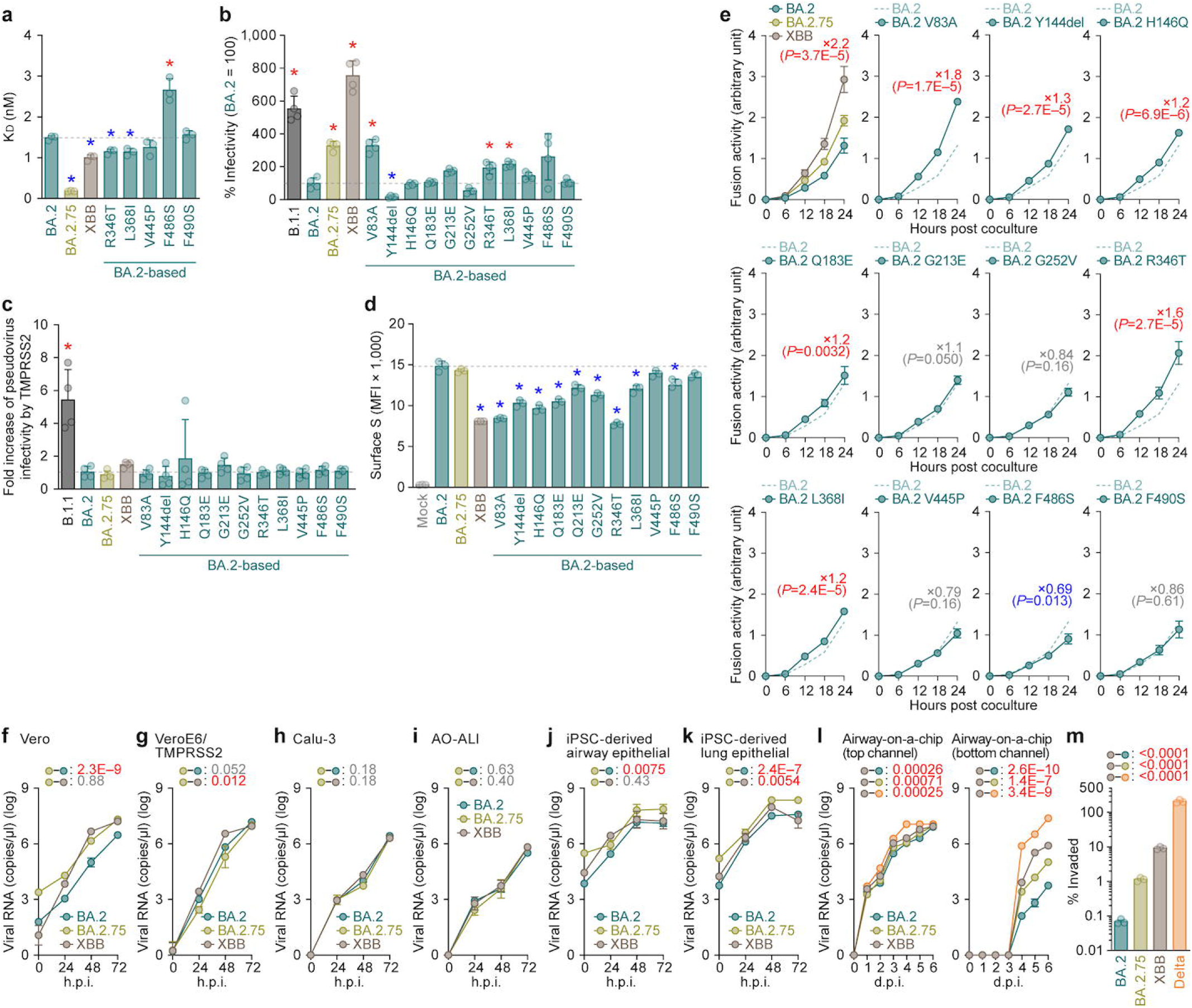
Virological characteristics of XBB *in vitro*. **a,** Binding affinity of the RBD of SARS-CoV-2 S protein to ACE2 by yeast surface display. The K_D_ value indicating the binding affinity of the RBD of the SARS-CoV-2 S protein to soluble ACE2 when expressed on yeast is shown. **b,** Pseudovirus assay. HOS-ACE2-TMPRSS2 cells were infected with pseudoviruses bearing each S protein. The amount of input virus was normalized based on the amount of HIV-1 p24 capsid protein. The percent infectivity compared to that of the virus pseudotyped with the BA.2 S protein are shown. **c,** Fold increase in pseudovirus infectivity based on TMPRSS2 expression. **d,e,** S-based fusion assay. **d,** S protein expression on the cell surface. The summarized data are shown. **e,** S-based fusion assay in Calu-3 cells. The recorded fusion activity (arbitrary units) is shown. The dashed green line indicates the result of BA.2. The red number in each panel indicates the fold difference between BA.2 and the derivative tested (XBB in the top left panel) at 24 h post coculture. **f–m,** Growth kinetics of XBB. Clinical isolates of BA.2, BA.2.75, XBB and Delta (only in **l,m**) were inoculated into Vero cells (**f**), VeroE6/TMPRSS2 cells (**g**), Calu-3 cells (**h**), AO-ALI (**i**), iPSC-derived airway epithelial cells (**j**), iPSC-derived lung epithelial cells (**k**) and an airway-on-a-chip system (**l**). The copy numbers of viral RNA in the culture supernatant (**f–h**), the apical sides of cultures (**i–k**), and the top (**l, left**) and bottom (**l, right**) channels of an airway-on-a-chip were routinely quantified by RT–qPCR. In **m**, the percentage of viral RNA load in the bottom channel per top channel at 6 d.p.i. (i.e., % invaded virus from the top channel to the bottom channel) is shown. Assays were performed in triplicate (**a,l,m**) or quadruplicate (**b–k**). The presented data are expressed as the average ± SD (**a–e**) or SEM (**f–m**). In **a–d,m**, each dot indicates the result of an individual replicate. In **a–d**, the dashed horizontal lines indicate the value of BA.2. In **a–d**, statistically significant differences (*, *P* < 0.05) versus BA.2 were determined by two-sided Student’s *t* tests. Red and blue asterisks, respectively, indicate increased and decreased values. In **e–l**, statistically significant differences versus BA.2.75 (**e–k**) or XBB (**l**) across timepoints were determined by multiple regression. In **m**, statistically significant differences versus XBB were determined by two-sided Student’s *t* tests. The FWERs calculated using the Holm method (**e–l**) or *P* values (**m**) are indicated in the figures.

We next assessed viral infectivity using pseudoviruses. As shown in **Fig. 3b**, the infectivity of XBB pseudovirus was 7.6-fold greater than that of BA.2 pseudovirus. Consistent with the results of yeast surface display assay (**Fig. 3a**), two substitutions in the RBD, R346T (1.9-fold) and L368I (2.2-fold), significantly increased pseudovirus infectivity (**Fig. 3b**). Additionally, although two substitutions in the NTD, Y144del (0.18-fold) and G252V (0.54-fold), significantly decreased pseudovirus infectivity, a substitution in the NTD, V83A (3.3-fold), significantly increased (**Fig. 3b**). Altogether, our results suggest that the XBB S augments its infectious potential through the multiple substitutions in the RBD (R346T, L368I and N460K) and NTD (V83A).

To assess the association of TMPRSS2 usage with the increased pseudovirus infectivity of XBB, we used HEK293-ACE2/TMPRSS2 cells and HEK293-ACE2 cells, on which endogenous surface TMPRSS2 is undetectable^23^, as target cells. As shown in **Fig. 3c**, the infectivity of XBB pseudovirus was not increased by TMPRSS2 expression, suggesting that TMPRSS2 is not associated with an increase in the infectivity of XBB pseudovirus.

### Fusogenicity of XBB S

The fusogenicity of XBB S was measured by the SARS-CoV-2 S-based fusion assay^2, 5, 10, 23–28^. We first assessed the fusogenicity of BA.2.75. Consistent with previous studies^10, 18^, the BA.2.75 S exhibited higher fusogenicity than the BA.2 S (**Extended Data Fig. 2a,b**). The assay using the BA.2 S derivatives that harbor respective BA.2.75-specific substitutions revealed that only the N460K substitution significantly increased fusogenicity (**Extended Data Fig. 2b**). We then assessed the fusogenicity of XBB S. As shown in **Fig. 3d**, the surface expression level of XBB was significantly lower than those of BA.2 and BA.2.75. The S-based fusion assay showed that the XBB S is significantly more fusogenic than BA.2 S (2.2-fold) and BA.2.75 S (1.5-fold) (**Fig. 3e**). To assess the determinant substitutions in the XBB S that are responsible for augmented fusogenicity, we used the BA.2 S-based derivatives that harbor respective XBB-specific substitutions. We revealed that particularly two substitutions, V83A and R346T, significantly increased fusogenicity (**Fig. 3e**). Together with the experiments focusing on BA.2.75 S (**Extended Data Fig. 2b**), our results suggest that two substitutions in the RBD (R346T and N460K) and a substitution in the NTD (V83A) contribute to the augmented fusogenicity of XBB S.

### Virological characteristics of XBB *in vitro*

To investigate the growth kinetics of XBB in *in vitro* cell culture systems, we inoculated clinical isolates of BA.2^23^, BA.2.75^10^, and XBB into multiple cell cultures. The growth kinetics of XBB in Vero cells (**Fig. 3f**), Calu-3 cells (**Fig. 3h**), human airway organoid-derived air-liquid interface (AO-ALI) system (**Fig. 3i**), and human induced pluripotent stem cell (iPSC)-derived airway epithelial cells (**Fig. 3j**) were comparable with those of BA.2.75. On the other hand, XBB more efficiently expanded in VeroE6/TMPRSS2 cells than BA.2.75 (**Fig. 3g**). Similar to our previous study^10^, the growth of BA.2.75 was significantly greater than that of BA.2 in human iPSC-derived alveolar epithelial cells (**Fig. 3k**). However, XBB was less replicative than BA.2.75 in this culture system (**Fig. 3k**).

To quantitatively assess the impact of XBB infection on the airway epithelial-endothelial barriers, we used an airway-on-a-chips system^2, 10, 29, 30^. By measuring the amount of virus that invades from the top channel (**Fig. 3l, left**) to the bottom channel (**Fig. 3l, right**), we are able to evaluate the ability of viruses to disrupt the airway epithelial-endothelial barriers. Notably, the percentage of virus that invaded the bottom channel of XBB-infected airway-on-chips was significantly higher than that of BA.2.75-infected airway-on-chips (**Fig. 3m**). Together with the findings of S-based fusion assay (**Fig. 3e**), these results suggest that XBB is higher fusogenic than BA.2.75.

### Virological characteristics of XBB *in vivo*

To investigate the virological features of XBB *in vivo*, we inoculated clinical isolates of Delta^26^, BA.2.75^10^, and XBB. Consistent with our previous studies^2, 10, 25, 26^, Delta infection resulted in weight loss (**Fig. 4a, left**). On the other hand, the body weights of BA.2.75- and XBB-infected hamsters were stable and comparable (**Fig. 4a, left**). We then analyzed the pulmonary function of infected hamsters as reflected by two parameters, enhanced pause (Penh) and the ratio of time to peak expiratory flow relative to the total expiratory time (Rpef). Among the four groups, Delta infection resulted in significant differences in these two respiratory parameters compared to XBB (**Fig. 4a, middle and right**), suggesting that XBB is less pathogenic than Delta. In contrast, although the Penh and Rpef values of XBB-infected hamsters were significantly different from those of uninfected hamsters, these were comparable to those of BA.2.75-infected hamsters (**Fig. 4a, middle and right**). These observations suggest that the pathogenicity of XBB is comparable to that of BA.2.75.

**Fig. 4.**
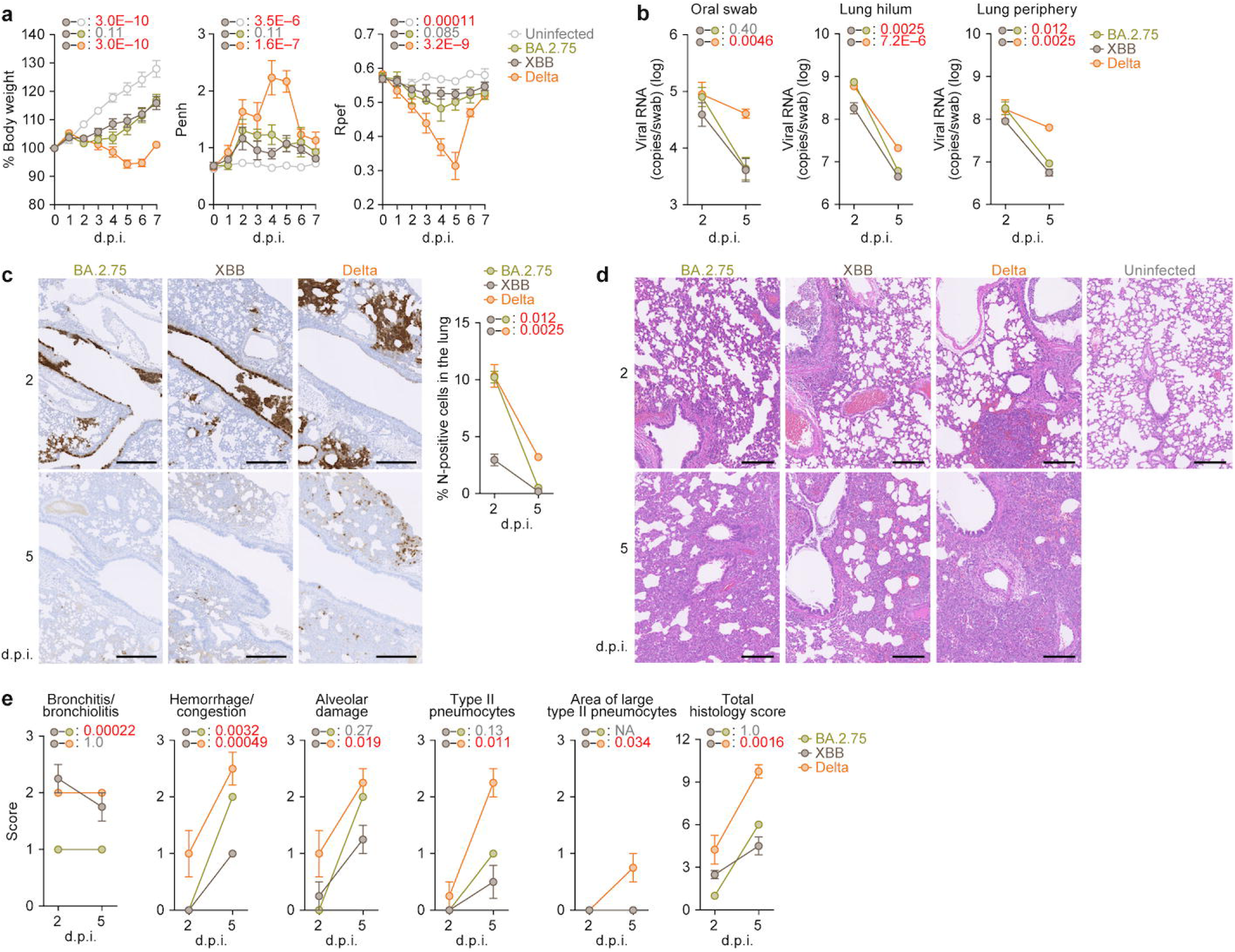
Virological characteristics of XBB *in vivo*. Syrian hamsters were intranasally inoculated with BA.2.75, XBB and Delta. Six hamsters of the same age were intranasally inoculated with saline (uninfected). Six hamsters per group were used to routinely measure the respective parameters (**a**). Four hamsters per group were euthanized at 2 and 5 d.p.i. and used for virological and pathological analysis (**b–e**). **a,** Body weight, Penh, and Rpef values of infected hamsters (n = 6 per infection group). **b,** (Left) Viral RNA loads in the oral swab (n=6 per infection group). (Middle and right) Viral RNA loads in the lung hilum (middle) and lung periphery (right) of infected hamsters (n=4 per infection group). **c,** IHC of the viral N protein in the lungs at 2 d.p.i. (top) and 5 d.p.i. (bottom) of infected hamsters. Representative figures (left, N-positive cells are shown in brown) and the percentage of N-positive cells in whole lung lobes (right, n=4 per infection group) are shown. The raw data are shown in **Extended Data Fig. 3. d,e,** H&E staining of the lungs of infected hamsters. Representative figures are shown in **d**. Uninfected lung alveolar space and bronchioles are also shown. **e,** Histopathological scoring of lung lesions (n=4 per infection group). Representative pathological features are reported in our previous studies^2,5,10,23,25,26^. In **a–c**, data are presented as the average ± SEM. In **a,b,c,e**, statistically significant differences between XBB and other variants across timepoints were determined by multiple regression. In **a**, the 0 d.p.i. data were excluded from the analyses. The FWERs calculated using the Holm method are indicated in the figures. Scale bars, 500 μm (**c**); 200 μm (**d**).

To address the viral spread in hamsters, we measured the viral RNA load in the oral swab. Although the viral RNA loads of the hamsters infected with XBB were significantly lower than those infected with Delta, there was no statistical difference between XBB and BA.2.75 (**Fig. 4b, left**). To assess the efficacy of viral spread in the respiratory tissues, we collected the lungs of infected hamsters at 2 and 5 d.p.i. and separated them into the hilum and periphery regions. However, the viral RNA loads in both lung hilum and periphery of XBB-infected hamsters were significantly lower than those of BA.2.75- and Delta-infected hamsters (**Fig. 4b, middle and right**), suggesting that XBB less efficiently spreads in the lungs of infected hamsters than BA.2.75 and XBB. We then investigated the viral spread in the respiratory tissues by immunohistochemical (IHC) analysis targeting viral nucleocapsid (N) protein. As shown in **Fig. 4c and Extended Data Fig. 3a**, the percentage of N-positive cells in the lungs of XBB-infected hamsters was significantly lower than those of BA.2.75- and Delta-infected hamsters. These data suggest that the spreading efficiency of XBB in the lungs of infected hamsters is comparable with or even lower than that of BA.2.75.

### Intrinsic pathogenicity of XBB

To investigate the pathogenicity of XBB in the lung, the formalin-fixed right lungs of infected hamsters were analyzed by carefully identifying the four lobules and main bronchus and lobar bronchi sectioning each lobe along with the bronchial branches. Histopathological scoring was performed as described in the previous studies^2, 5, 10, 23, 25, 26^. Briefly, bronchitis or bronchiolitis, hemorrhage or congestion, alveolar damage with epithelial apoptosis and macrophage infiltration, type II pneumocytes and the area of the hyperplasia of large type II pneumocytes were evaluated by certified pathologists and the degree of these pathological findings were arbitrarily scored using a four-tiered system as 0 (negative), 1 (weak), 2 (moderate), and 3 (severe)^2, 5, 10, 23, 25, 26^. Similar to our previous studies^2, 10, 25, 26^, four out of the five histological parameters as well as the total score of Delta-infected hamsters were significantly greater than those of XBB-infected hamsters (**Fig. 4e**). When we compared the histopathological scores of two Omicron subvariants, the scores of type II pneumocytes and the area of the hyperplasia of large type II pneumocytes, and total histology score of XBB-infected hamsters were comparable with those of BA.2.75-infected hamsters (**Fig. 4e**). Altogether, these histopathological analyses suggest that the intrinsic pathogenicity of XBB is lower than that of Delta and comparable with that of BA.2.75.

## Discussion

Here, we illuminated the evolutionary and epidemic dynamics of XBB variant, a recombinant lineage rapidly spreading around the world. Our phylogenetic analyses suggested that XBB emerged through the recombination of two co-circulating BA.2 lineages, BJ.1 and BM.1.1.1 (a progeny of BA.2.75), during the summer of 2022 in India or its neighboring countries (**Fig. 1**). Furthermore, XBB shows substantially higher R_e_ than the parental lineages, suggesting that the recombination event increased R_e_ (i.e., viral fitness). To our knowledge, this is the first documented example of a SARS-CoV-2 variant increasing its fitness through recombination rather than single mutations. Furthermore, we showed that the R_e_ values of XBB lineages are comparable with or slightly higher than those of BQ.1 lineages, and XBB and BQ.1 lineages are becoming dominants in Eastern and Western Hemisphere, respectively. Since XBB lineages are rapidly spreading also in Western countries such as the USA, this regional difference is likely explained simply by the distance from the emergence regions of these lineages rather than caused by the fact that viral fitness changes depending on circulated regions. Together, although XBB now circulates mainly in countries around India, this variant has a potential to spread worldwide in the near future.

Compared to BA.5, BA.2.75 and even BQ.1.1 (ref.^2^), the most remarkable feature of XBB is the profound resistance to antiviral humoral immunity induced by breakthrough infections of prior Omicron subvariants (**Fig. 2**). In fact, our analyses showed that 10 out of 14 breakthrough BA.2 infection sera and 9 out of 20 breakthrough BA.5 infection sera fail to neutralize XBB. The neutralization experiments using single mutants showed that multiple substitutions in the XBB S protein cooperatively contribute to the immune resistance of XBB, and particularly, not only the substitutions in the RBD but also at least a mutation in the NTD, Y144del, closely associates with the immune resistant property of XBB. The effect of Y144del mutation on immune resistance is reported in a recent study by Cao et al.^12^, and this mutation has been observed in previous variants of concern such as Alpha and Omicron BA.1. Furthermore, we previously showed that the Mu variant, one of the previous variants of interest, also has a mutation in the region including Y144 (i.e., YY144-145TSN), which contributes to the robust immune escape of this variant^31, 32^. Additionally, the region including Y144 is proposed that it is one of the major epitopes of NTD targeting neutralization antibodies^33^. Together, these observations suggest that Y144del mutation in XBB S contributed to escape from these NTD targeting neutralization antibodies.

A series of our previous studies^2, 5, 10^ showed that the substitutions in S proteins associating with the escape from humoral immunity tend to lead to acquire substitutions enhancing the ACE2 binding affinity or viral infectivity probably to compensate for the negative effects of the immune escape-associated substitutions on the ACE2 binding affinity. In the present study, we show that XBB harbors both the immune escape-associated substitutions (i.e., Y144del and F486S) and the infectivity-enhancing substitutions (i.e., V83A and N460K). Importantly, XBB emerged through recombination in the *S* gene, and Y144del and V83A are located on the 5’ recombinant fragment while F486S and N460K are on the 3’ fragment. This means that XBB acquired two sets of a pair of immune escape-associated and infectivity-enhancing substitutions by only one recombination event. Harboring the two sets of the substitution pairs would be one of the causes why XBB shows higher R_e_ than other Omicron subvariants. Together, although XBB emerged via a unique evolutionary pathway, our data suggest that XBB also follows the same evolutionary rule with other Omicron subvariants.

Although BQ.1.1, another Omicron subvariant of concern at the end of 2022, acquired these two types of mutation by convergent substitutions^2^, XBB acquired them by a recombination event. In terms of the evolutionary strategy to acquire these two types of mutations, XBB emerged in a unique way.

In our previous studies focusing on some Omicron subvariants such as BA.1 (ref.^25^), BA.2 (ref.^23^), BA.5 (ref.^5^), and BA.2.75 (ref.^10^), viral fusogenicity in *in vitro* experimental setup was well correlated to viral intrinsic pathogenicity in a hamster model. However, although the fusogenicity of XBB was greater than that of BA.2.75, one of the parental lineages of XBB, the intrinsic pathogenicity of XBB was comparable or even lower than that of BA.2.75. The discrepancy between viral fusogenicity and intrinsic pathogenicity was also observed in another Omicron subvariant of concern at the end of 2022, BQ.1.1 (ref.^2^). The discrepancy between viral fusogenicity and intrinsic pathogenicity may be explained by at least three possibilities. First, certain mutations in the non-S region of the XBB genome can attenuate viral pathogenicity augmented by the higher fusogenicity compared with BA.2.75. There are at least seven substitutions in the non-S region of XBB when compared to that of BA.2.75, and some of these mutations may attenuate the viral intrinsic pathogenicity (**Extended Data Fig. 1a**). Second, a theoretical study by Sasaki, Lion, and Boots provided a possibility that antigenic escape can augment viral pathogenicity^34^. Since we demonstrated that at least two descendants of BA.2, BA.5 (ref.^5^) and BA.2.75 (ref.^10^), increased their intrinsic pathogenicity, this theory may explain the evolution of Omicron. More importantly, this theory also predicts that there is a limitation to increase viral pathogenicity^34^. Together with our observations, it might be possible to assume that the pathogenicity of Omicron lineage already reaches a plateau. Third, in the cases of BQ.1.1 (ref.^2^) and XBB, it might be possible that the tropism and affinity of S proteins of these variants are different between human ACE2 and hamster ACE2, and therefore, a hamster model may not reproduce the human condition.

In summary, our results suggested that XBB is highly transmissible and resistant to the antiviral humoral immunity induced by breakthrough infections of continued in-depth viral genomic surveillance and real-time evaluation of the risk of newly emerging SARS-CoV-2 variants, even though considered local variants at the time of emergence, should be crucial.

## Author Contributions

Jumpei Ito performed bioinformatics, modeling, and statistical analysis. Jumpei Ito and Spyros Lytras performed phylogenetic analyses. Keiya Uriu, Hesham Nasser, Maya Shofa, Sayaka Deguchi, MST Monira Begum, Yusuke Kosugi, Shigeru Fujita, Terumasa Ikeda, Akatsuki Saito and Kazuo Takayama performed cell culture experiments.

Tomokazu Tamura, Izumi Kida, Naganori Nao, Yukari Itakura, Rigel Suzuki, Kumiko Yoshimatsu, Saori Suzuki, Takasuke Fukuhara and Keita Matsuno performed animal experiments.

Yoshitaka Oda, Lei Wang, Masumi Tsuda and Shinya Tanaka performed histopathological analysis.

Jiri Zahradnik and Gideon Schreiber performed yeast surface display assay. Sayaka Deguchi and Kazuo Takayama prepared AO, AO-ALI and airway-on-a-chip systems.

Yuki Yamamoto and Tetsuharu Nagamoto performed generation and provision of human iPSC-derived airway and alveolar epithelial cells.

Hiroyuki Asakura, Mami Nagashima, Kenji Sadamasu and Kazuhisa Yoshimura performed viral genome sequencing analysis.

Jumpei Ito performed statistical, modelling, and bioinformatics analyses.

Jumpei Ito, Terumasa Ikeda, Takasuke Fukuhara, Akatsuki Saito, Shinya Tanaka, Keita Matsuno, Kazuo Takayama and Kei Sato designed the experiments and interpreted the results.

Jumpei Ito and Kei Sato wrote the original manuscript. All authors reviewed and proofread the manuscript.

The Genotype to Phenotype Japan (G2P-Japan) Consortium contributed to the project administration.

## Acknowledgments

We would like to thank all members belonging to The Genotype to Phenotype Japan (G2P-Japan) Consortium. We thank Dr. Kenzo Tokunaga (National Institute for Infectious Diseases, Japan) and Dr. Jin Gohda (The University of Tokyo, Japan) for providing reagents. We also thank National Institute for Infectious Diseases, Japan for providing clinical isolates of BQ.1.1 (strain TY41-796-P1; GISAID ID: EPI_ISL_15579783) and BA.2 (strain TY40-385; GISAID ID: EPI_ISL_9595859). We appreciate the technical assistance from The Research Support Center, Research Center for Human Disease Modeling, Kyushu University Graduate School of Medical Sciences. We gratefully acknowledge all data contributors, i.e. the Authors and their Originating laboratories responsible for obtaining the specimens, and their Submitting laboratories for generating the genetic sequence and metadata and sharing via the GISAID Initiative, on which this research is based. The super-computing resource was provided by Human Genome Center at The University of Tokyo.

This study was supported in part by AMED SCARDA Japan Initiative for World-leading Vaccine Research and Development Centers “UTOPIA” (JP223fa627001, to Kei Sato), AMED SCARDA Program on R&D of new generation vaccine including new modality application (JP223fa727002, to Kei Sato); AMED Research Program on Emerging and Re-emerging Infectious Diseases (JP21fk0108574, to Hesham Nasser; JP21fk0108481, to Akatsuki Saito; JP21fk0108465, to Akatsuki Saito; JP21fk0108493, to Takasuke Fukuhara; JP22fk0108617 to Takasuke Fukuhara; JP22fk0108146, to Kei Sato; JP21fk0108494 to G2P-Japan Consortium, Keita Matsuno, Shinya Tanaka, Terumasa Ikeda, Takasuke Fukuhara, and Kei Sato; JP21fk0108425, to Kazuo Takayama, Akatsuki Saito and Kei Sato; JP21fk0108432, to Kazuo Takayama, Takasuke Fukuhara and Kei Sato); AMED Research Program on HIV/AIDS (JP22fk0410033, to Akatsuki Saito; JP22fk0410047, to Akatsuki Saito; JP22fk0410055, to Terumasa Ikeda; and JP22fk0410039, to Kei Sato); AMED CRDF Global Grant (JP22jk0210039 to Akatsuki Saito); AMED Japan Program for Infectious Diseases Research and Infrastructure (JP22wm0325009, to Akatsuki Saito; JP22wm0125008 to Keita Matsuno); AMED CREST (JP21gm1610005, to Kazuo Takayama); JST PRESTO (JPMJPR22R1, to Jumpei Ito); JST CREST (JPMJCR20H4, to Kei Sato); JSPS KAKENHI Grant-in-Aid for Scientific Research C (22K07103, to Terumasa Ikeda); JSPS KAKENHI Grant-in-Aid for Scientific Research B (21H02736, to Takasuke Fukuhara); JSPS KAKENHI Grant-in-Aid for Early-Career Scientists (22K16375, to Hesham Nasser; 20K15767, Jumpei Ito); JSPS Core-to-Core Program (A. Advanced Research Networks) (JPJSCCA20190008, Kei Sato); JSPS Research Fellow DC2 (22J11578, to Keiya Uriu); JSPS Leading Initiative for Excellent Young Researchers (LEADER) (to Terumasa Ikeda); World-leading Innovative and Smart Education (WISE) Program 1801 from the Ministry of Education, Culture, Sports, Science and Technology (MEXT) (to Naganori Nao); The Tokyo Biochemical Research Foundation (to Kei Sato); Takeda Science Foundation (to Terumasa Ikeda); Mochida Memorial Foundation for Medical and Pharmaceutical Research (to Terumasa Ikeda); The Naito Foundation (to Terumasa Ikeda); Shin-Nihon Foundation of Advanced Medical Research (to Terumasa Ikeda); Waksman Foundation of Japan (to Terumasa Ikeda); an intramural grant from Kumamoto University COVID-19 Research Projects (AMABIE) (to Terumasa Ikeda); Ito Foundation Research Grant R4 (to Akatsuki Saito); and the project of National Institute of Virology and Bacteriology, Programme EXCELES, funded by the European Union, Next Generation EU (LX22NPO5103, to Jiri Zahradnik).

## Consortia

Hirofumi Sawa^7, 13, 14^, Marie Kato^11^, Zannatul Ferdous^11^, Hiromi Mouri^11^, Kenji Shishido^11^, Naoko Misawa^2^, Izumi Kimura^2^, Lin Pan^2^, Mai Suganami^2^, Mika Chiba^2^, Ryo Yoshimura^2^, Kyoko Yasuda^2^, Keiko Iida^2^, Naomi Ohsumi^2^, Adam P. Strange^2^, Daniel Sauter^2,30^, So Nakagawa^31^ Jiaqi Wu^31^, Rina Hashimoto^16^, Yukio Watanabe^16^, Ayaka Sakamoto^16^, Naoko Yasuhara^16^, Takao Hashiguchi^32^, Tateki Suzuki^32^, Kanako Kimura^32^, Jiei Sasaki^32^, Yukari Nakajima^32^, Hisano Yajima^32^, Kotaro Shirakawa^32^, Akifumi Takaori-Kondo^32^, Kayoko Nagata^32^, Yasuhiro Kazuma^32^, Ryosuke Nomura^32^, Yoshihito Horisawa^32^, Yusuke Tashiro^32^, Yugo Kawa^32^, Takashi Irie^33^, Ryoko Kawabata^33^, Ryo Shimizu^7^, Otowa Takahashi^7^, Kimiko Ichihara^7^, Chihiro Motozono^34^, Mako Toyoda^34^, Takamasa Ueno^34^, Yuki Shibatani^9^, Tomoko Nishiuchi^9^

^30^University Hospital Tübingen, Tübingen, Germany ^31^Tokai University School of Medicine, Isehara, Japan ^32^Kyoto University, Kyoto, Japan

^33^Hiroshima University, Hiroshima, Japan

^34^Kumamoto University, Kumamoto, Japan

## Figure legends

**Supplementary Table 1**. Estimated relative R_e_ values of XBB-related lineages in India

**Supplementary Table 2.** Estimated lineage frequencies for BQ.1, XBB other lineages as of November 15, 2022 in each country

**Supplementary Table 3.** Estimated relative R_e_ values of viral lineages by a hierarchical Bayesian model

**Supplementary Tables 4.** Human sera used in this study

**Supplementary Table 5.** Primers used for the construction of SARS-CoV-2 S expression plasmids

**Supplementary Table 6.** Summary of unexpected amino acid mutations detected in the working virus stocks

**Extended Data Fig. 1. Phylogenetic and epidemic analyses of the XBB lineage**

**a,** Ancestral state reconstruction of the circulated regions of viruses, related to **Fig. 1e**. Unlike **Fig. 1e**, result for the entire of the tree is shown. An asterisk denotes the MRCA node of XBB lineages.

**b**, Relative R_e_ values for viral lineages in India, assuming a fixed generation time of 2.1 days, related to **Fig. 1f**. The R_e_ of BA.2 is set at 1. The posterior (violin), posterior mean (dot), and 95% Bayesian CI (line) are shown.

**c,** Relative R_e_ values for viral lineages, assuming a fixed generation time of 2.1 days, related to **Fig. 1h**. The R_e_ value of BA.5 is set at 1. The posterior (violin), posterior mean (dot), and 95% Bayesian CI (line) are shown. R_e_ values for each country where BQ.1 and XBB lineages cocirculated are shown.

**Extended Data Fig. 2. Fusogenicity of BA.2.75 S**

**a,b,** S-based fusion assay. **a,** S protein expression on the cell surface. The summarized data are shown. **b,** S-based fusion assay in Calu-3 cells. The recorded fusion activity (arbitrary units) is shown. The dashed green line indicates the result of BA.2. The red number in each panel indicates the fold difference between BA.2 and the derivative tested (XBB in the top left panel) at 24 h post coculture.

Assays were performed in triplicate. The presented data are expressed as the average ± SD. In **a**, each dot indicates the result of an individual replicate, and the dashed horizontal lines indicate the value of BA.2. Statistically significant differences (*, *P* < 0.05) versus BA.2 were determined by two-sided Student’s *t* tests, and red asterisks indicate increased values. In **b**, statistically significant differences versus BA.2 across timepoints were determined by multiple regression. The FWERs calculated using the Holm method are indicated in the figures.

**Extended Data Fig. 3. Histological observations in infected hamsters**

IHC of the SARS-CoV-2 N protein in the lungs of infected hamsters at 2 d.p.i. (**a**) and 5 d.p.i (**b**) (4 hamsters per infection group). In each panel, IHC staining (top) and the digitalized N-positive area (bottom, indicated in red) are shown. The red numbers in the bottom panels indicate the percentage of the N-positive area. Summarized data are shown in **Fig. 4c, right**. Scale bars, 5 mm.

## Methods

### Ethics statement

All experiments with hamsters were performed in accordance with the Science Council of Japan’s Guidelines for the Proper Conduct of Animal Experiments. The protocols were approved by the Institutional Animal Care and Use Committee of National University Corporation Hokkaido University (approval ID: 20-0123 and 20-0060). All protocols involving specimens from human subjects recruited at Interpark Kuramochi Clinic was reviewed and approved by the Institutional Review Board of Interpark Kuramochi Clinic (approval ID: G2021-004). All human subjects provided written informed consent. All protocols for the use of human specimens were reviewed and approved by the Institutional Review Boards of The Institute of Medical Science, The University of Tokyo (approval IDs: 2021-1-0416 and 2021-18-0617) and University of Miyazaki (approval ID: O-1021).

### Human serum collection

Convalescent sera were collected from fully vaccinated individuals who had been infected with BA.2 (9 2-dose vaccinated and 5 3-dose vaccinated; 11–61 days after testing. n=14 in total; average age: 47 years, range: 24–84 years, 64% male) (**Fig. 2a**), and fully vaccinated individuals who had been infected with BA.5 (2 2-dose vaccinated, 17 3-dose vaccinated and 1 4-dose vaccinated; 10–23 days after testing. n=20 in total; average age: 51 years, range: 25–73 years, 45% male) (**Fig. 2b**). The SARS-CoV-2 variants were identified as previously described^5,10, 23^. Sera were inactivated at 56°C for 30 minutes and stored at –80°C until use. The details of the convalescent sera are summarized **in** Supplementary Table 4.

### Cell culture

HEK293T cells (a human embryonic kidney cell line; ATCC, CRL-3216), HEK293 cells (a human embryonic kidney cell line; ATCC, CRL-1573) and HOS-ACE2/TMPRSS2 cells (HOS cells stably expressing human ACE2 and TMPRSS2)^35, 36^ were maintained in DMEM (high glucose) (Sigma-Aldrich, Cat# 6429-500ML) containing 10% fetal bovine serum (FBS, Sigma-Aldrich Cat# 172012-500ML) and 1% penicillin–streptomycin (PS) (Sigma-Aldrich, Cat# P4333-100ML). HEK293-ACE2 cells (HEK293 cells stably expressing human ACE2)^24^ were maintained in DMEM (high glucose) containing 10% FBS, 1 µg/ml puromycin (InvivoGen, Cat# ant-pr-1) and 1% PS. HEK293-ACE2/TMPRSS2 cells (HEK293 cells stably expressing human ACE2 and TMPRSS2)^24^ were maintained in DMEM (high glucose) containing 10% FBS, 1 µg/ml puromycin, 200 µg/ml hygromycin (Nacalai Tesque, Cat# 09287-84) and 1% PS. Vero cells [an African green monkey (*Chlorocebus sabaeus*) kidney cell line; JCRB Cell Bank, JCRB0111] were maintained in Eagle’s minimum essential medium (EMEM) (Sigma-Aldrich, Cat# M4655-500ML) containing 10% FBS and 1% PS. VeroE6/TMPRSS2 cells (VeroE6 cells stably expressing human TMPRSS2; JCRB Cell Bank, JCRB1819)^37^ were maintained in DMEM (low glucose) (Wako, Cat# 041-29775) containing 10% FBS, G418 (1 mg/ml; Nacalai Tesque, Cat# G8168-10ML) and 1% PS. Calu-3 cells (ATCC, HTB-55) were maintained in Eagle’s minimum essential medium (EMEM) (Sigma-Aldrich, Cat# M4655-500ML) containing 10% FBS and 1% PS. Calu-3/DSP_1-7_ cells (Calu-3 cells stably expressing DSP_1-7_)^38^ were maintained in EMEM (Wako, Cat# 056-08385) containing 20% FBS and 1% PS. Human airway and lung epithelial cells derived from human induced pluripotent stem cells (iPSCs) were manufactured according to established protocols as described below (see “Preparation of human airway and lung epithelial cells from human iPSCs” section) and provided by HiLung Inc. AO-ALI model was generated according to established protocols as described below (see “AO-ALI model” section).

### Viral genome sequencing

Viral genome sequencing was performed as previously described^5^. Briefly, the virus sequences were verified by viral RNA-sequencing analysis. Viral RNA was extracted using a QIAamp viral RNA mini kit (Qiagen, Cat# 52906). The sequencing library employed for total RNA sequencing was prepared using the NEBNext Ultra RNA Library Prep Kit for Illumina (New England Biolabs, Cat# E7530). Paired-end 76-bp sequencing was performed using a MiSeq system (Illumina) with MiSeq reagent kit v3 (Illumina, Cat# MS-102-3001). Sequencing reads were trimmed using fastp v0.21.0 (ref.^39^) and subsequently mapped to the viral genome sequences of a lineage B isolate (strain Wuhan-Hu-1; GenBank accession number: NC_045512.2)^37^ using BWA-MEM v0.7.17 (ref.^40^). Variant calling, filtering, and annotation were performed using SAMtools v1.9 (ref.^41^) and snpEff v5.0e^42^.

### Recombination analysis

As of October 3, 2022, we retrieved a total of 562 sequences satisfying the following criteria from the GISAID database (https://gisaid.org/): i) human hosts, ii) collected after 2022, iii) with length greater than 28,000 base pairs, and iv) with PANGO lineage designation BJ.1, BM.1, XBB and all their descendants. To ensure that PANGO lineage definitions in our dataset’s metadata included the latest circulating lineages, the GISAID metadata were downloaded again on October 15, 2022, and the PANGO lineages of our sequences were updated accordingly. Sequences were aligned to the reference Wuhan-Hu-1 genome (GenBank Accession no. NC_045512.2) and then converted to a multiple sequence alignment using the ‘global_profile_alignment.sh’ script from the SARS-CoV-2 global phylogeny pipeline^43^, utilizing MAFFT^44^. A number of recombination detection methods were performed on the resulting alignment using the Recombination Detection Program (RDP) v.5.21 (ref.^19^), specifically: RDP^45^, GENECONV^46^, Chimaera^47^, MaxChi^48^, 3seq^49^, BootScan^50^ and SiScan^51^. Sequences were assumed to be linear, only recombination events detected consistently by more than 3 independent methods were retrieved and potential false positives were excluded from the final output of RDP5.

### Phylogenetic analyses

To reconstruct the overall relatedness of the XBB parent lineages BJ.1 and BM.1.1.1 to the other Omicron variants (**Fig. 1a**) we retrieved 100 random sequences from each Omicron PANGO lineages: BA.1, BA.2, BA.4 and BA.5 and 20 random sequences from each younger lineage: BQ.1.1, BA.2.75, BJ.1, and BM.1.1.1. Sequence EPI_ISL_466615 was also added as an outgroup, representing the oldest isolate of B.1.1 obtained in the UK. The sequences were aligned to the reference Wuhan-Hu-1 genome (NC_045512.2) and then converted to a multiple sequence alignment using the ‘global_profile_alignment.sh’ script from the SARS-CoV-2 global phylogeny pipeline^43^ utilizing MAFFT^44^. Fasttree v.2.1 (ref.^52^) was used to infer the phylogeny for the nucleotide alignment under a GTR substitution model (option -gtr).

For inferring the phylogenies of each non-recombinant segment of the XBB variant, we first split the alignment used for the recombination analysis above at genome position 22,920 (the breakpoint inferred by RDP5). Due to the lack of many informative sites of the 3’ end shorter non-recombinant alignment, two quality filtering steps were implemented: i) the 3’ end of the alignment was trimmed up to the position where none of the sequences had 3’ end gaps and ii) all sequences with Ns were removed, leading to a reduced alignment of 370 sequences. BA.2 sequence EPI_ISL_10926749 was added to the alignments as an outgroup. Iqtree2 v2.1.3 (ref.^53^) was used for making a phylogenetic for each non-recombinant alignment. The TIM2+F+I substitution model was used for both trees as selected by the ‘-m TEST’ of iqtree and node support was assessed by performing 1000 ultrafast bootstrap replicates.

Both phylogenies were manually inspected for the presence of temporal signal using TempEst v1.5.3 (ref.^54^). The 3’ end non-recombinant segment’s phylogeny did not have enough substitutions for a root-to-tip regression to be inferred, hence we proceeded with tip-dating analysis only for the 5’ end, longer segment. The 5’ end phylogeny was manually rooted to the BA.2 outgroup branch with FigTree [https://github.com/rambaut/figtree/] and TreeTime v.0.8.1 (ref.^55^) was used for the time-calibration (maintaining the rooting with the –keep-root option). The substitution rate inferred by the root-to-tip regression and used for the tree calibration was 1.10E-3 substitutions per site per year, consistent with the accepted rate for SARS-CoV-2 (ref.^56^). Two terminal branches with dates not matching the root-to-tip regression were manually removed from the phylogeny.

To infer the emergence region of XBB, we performed a discrete ancestral state reconstruction analysis utilizing a Markov model. In this analysis, the time-calibrated phylogenetic tree of the 5’ non-recombinant segment (1–22,920) (shown in **Fig. 1d, left**) was used. First, based on the sampled (or circulating) region of each virus in the tree, we defined the discrete state of the circulating region for each viral sequence according to the following rules: For samples from India, Bangladesh, and Singapore, the states “India”, “Bangladesh”, and “Singapore” were assigned respectively. For samples from the other Asian countries, “Other Asian countries” was assigned. For samples from European and Northern American countries, “European countries” and “Northern American countries” were assigned, respectively. For samples from the other regions, “Other regions” was assigned. We subsequently inferred the ancestral likelihoods of the seven states (i.e., circulating regions) for each internal node of the tree using the asr_mk_model function in the castor package^57^. In the tree shown in **Fig. 1e and Extended Data Fig. 1b**, we mapped the ancestral likelihood of the state “India” to the tree. The analyses above were performed on R v.4.2.1.

### Epidemic dynamics analyses

We modeled the epidemic dynamics of viral lineages based on the viral genomic surveillance data deposited in the GISAID database (https://www.gisaid.org/). In the present study, we performed three types of analyses: i) The estimation of the relative R_e_ for lineages related to XBB in India (shown in **Fig. 1f and Extended Data Fig. 1c**), ii) The estimation of the epidemic frequencies of XBB and BQ.1 lineages in each country as of November 15, 2022 (shown in **Fig. 1g**), and iii) The estimation of the global and country-specific R_e_ value of XBB and BQ.1 lineages in the countries where these variants circulated (**Fig. 1h,i and Extended Data Fig. 1d**). For the three analyses, the metadata of viral sequences downloaded from the GISAID database on December 1^st^, 2022 was used. We excluded the sequence records with the following features: i) a lack of collection date information; ii) sampling in animals other than humans; iii) sampling by quarantine; or iv) without the PANGO lineage information.

To estimate the relative R_e_ for lineages related to XBB in India, we analyzed the records for samples from India from June 1, 2022 to November 15, 2022. We removed records with >5% undetermined (N) nucleotide sequences from the dataset. We first simplified the viral lineage classification based on the PANGO lineage. We renamed the sublineages of BA.5 as BA.5, and subsequently, we removed the BA.5 sequences harboring any of the convergent S substitutions, S:R346X, S:K444X, and S:N460X from our dataset in order to exclude the sequences belonging to the recent BA.5 sublineages exhibiting particularly higher R_e_ such as BQ.1.1 (ref.^2^). Also, we removed the sequences of BA.2.75 harboring any of the convergent S substitutions, S:R346X, S:K444X, S:N460X, and S:F486X. Furthermore, since a part of BA.2.10.1 sequences harbor XBB-characteristic substitutions (S:V83A, S:F486S, and S:F490S) probably due to the misclassification of XBB, we removed the sequences of BA.2.10.1 harboring these XBB-characteristic substitutions. According to the modified viral lineages, we extracted records for viral lineages of interest: BA.2, BA.5, BA.2.75, BM.1, BM1.1, BM.1.1.1, BA.2.10, BJ.1, XBB, and XBB.1. Subsequently, we counted the daily frequency of each viral lineage. Relative R_e_ value for each viral lineage was estimated according to the Bayesian multinomial logistic model, described in our previous study^5^. Briefly, we estimated the logistic slope parameter for each viral lineage using the model and then calculated relative R_e_ for each lineage *r_l_* as *r_l_* = *exp*(γβ_*l*_) where γ is the average viral generation time (2.1 days) (http://sonorouschocolate.com/covid19/index.php?title=Estimating_Generation_Time_Of_Omicron). Parameter estimation was performed via the MCMC approach implemented in CmdStan v2.30.1 (https://mc-stan.org) with CmdStanr v0.5.3 (https://mc-stan.org/cmdstanr/). Four independent MCMC chains were run with 500 and 1,000 steps in the warmup and sampling iterations, respectively. We confirmed that all estimated parameters showed <1.01 R-hat convergence diagnostic values and >200 effective sampling size values, indicating that the MCMC runs were successfully convergent. Information on the estimated parameters is summarized in **Supplementary Table 1**.

To estimate the epidemic frequencies of XBB and BQ.1 lineages in each country as of November 15, 2022, we analyzed the records for viral samples collected from August 1, 2022 to November 15^th^, 2022. In data for each country, we counted the daily lineage frequency of BQ.1 (including its decedent sublineages), XBB (including its decedent sublineages), and the other SARS-CoV-2 lineages (referred to as “Other lineages”). We analyzed the data ≥50 samples of either the BQ.1 or XBB lineages. In this criterion, 56 countries remained. Subsequently, we fitted the multinomial logistic model described in the paragraph above to the daily lineage frequency data of each country separately, and the epidemic frequency of each viral lineage as of November 15, 2022 in each country was estimated. If the data for November 15, 2022 in a particular country are not available, the lineage frequencies at the latest date in the country were used instead. The estimated lineage frequencies for BQ.1 and XBB in each country were shown on the global map using The R library maps v3.4.1 (https://cran.r-project.org/web/packages/maps/index.html). Information on the estimated lineage frequencies is summarized in **Supplementary Table 2**.

To estimate the global average and country-specific R_e_ values for BQ.1 and XBB lineages, we analyzed the sequence records for viral samples collected from August 1, 2022 to November 15, 2022. We defined the sequences of BQ.1 (including its sublineages) harboring S:R346T as BQ.1.1 and the other BQ.1 sequences as BQ.1. Similarly, the sequences of XBB (including its sublineages) harboring S:G252V as XBB and the other XBB sequences as XBB. Subsequently, we extracted the sequence records of BQ.1, BQ.1.1, XBB, and XBB.1 in addition to BA.5 (including its sublineages) and BA.2.75 (including its sublineages), which are predominant lineages before the BQ.1 and XBB emergencies. Next, we counted the daily frequency of the lineages above in 1000 samples and ≥200 samples of either the BQ.1, BQ.1.1, XBB, or XBB.1 lineages. In this criterion, 11 countries (Australia, Austria, Denmark, India, Indonesia, Israel, Malaysia, Peru, Singapore, the UK, and the USA) remained. To estimate the global average R_e_ values of the lineages above, we employed a hierarchal Bayesian multinomial logistic model, which we established in our previous studies^10, 23^. Briefly, this hierarchal model can estimate the global average and country-specific R_e_ values of lineages of interest simultaneously according to the daily lineage frequency data from multiple countries. The relative R_e_ of each viral lineage \ in each county (^i.^) was calculated according to the country-specific slope parameter, β_*ls*_, as r_*ls*_ = *exp*(γβ_*ls*_) where 1. is the average viral generation time (2.1 days). Similarly, the global average relative R_e_ of each viral lineage was calculated according to the global average slope parameter, β_*l*_, as r_*l*_ = *exp*(γβ_*ls*_). For parameter estimation, the global average intercept and slope parameters of the BA.5 variant were fixed at 0. Consequently, the relative R_e_ of BA.5 was fixed at 1, and those of the other lineages were estimated relative to that of BA.5. Parameter estimation was performed via the MCMC approach implemented in CmdStan v2.30.1 (https://mc-stan.org) with CmdStanr v0.5.3 (https://mc-stan.org/cmdstanr/). Four independent MCMC chains were run with 500 and 2,000 steps in the warmup and sampling iterations, respectively. We confirmed that all estimated parameters showed <1.01 R-hat convergence diagnostic values and >200 effective sampling size values, indicating that the MCMC runs were successfully convergent. Information on the estimated parameters is summarized in **Supplementary Table 3**.

### Plasmid construction

Plasmids expressing the codon-optimized SARS-CoV-2 S proteins of B.1.1 (the parental D614G-bearing variant), BA.2 and BA.5, BQ.1.1 and BA.2.75 were prepared in our previous studies^2, 5, 10, 23, 24, 58^. Plasmids expressing the codon-optimized S proteins of XBB and BA.2 S-based derivatives were generated by site-directed overlap extension PCR using the primers listed in **Supplementary Table 5**. The resulting PCR fragment was digested with KpnI (New England Biolabs, Cat# R0142S) and NotI (New England Biolabs, Cat# R1089S) and inserted into the corresponding site of the pCAGGS vector^59^. Nucleotide sequences were determined by DNA sequencing services (Eurofins), and the sequence data were analyzed by Sequencher v5.1 software (Gene Codes Corporation).

### Neutralization assay

Pseudoviruses were prepared as previously described^2, 5, 10, 23, 26, 28, 31, 32, 36, 38, 58, 60^. Briefly, lentivirus (HIV-1)-based, luciferase-expressing reporter viruses were pseudotyped with SARS-CoV-2 S proteins. HEK293T cells (1,000,000 cells) were cotransfected with 1 μg psPAX2-IN/HiBiT^35^, 1 μg pWPI-Luc2^35^, and 500 ng plasmids expressing parental S or its derivatives using PEI Max (Polysciences, Cat# 24765-1) according to the manufacturer’s protocol. Two days posttransfection, the culture supernatants were harvested and centrifuged. The pseudoviruses were stored at –80°C until use.

The neutralization assay (**Fig. 2**) was prepared as previously described^2, 5, 10, 23, 26, 28, 31, 32, 36, 38, 58, 60^. Briefly, the SARS-CoV-2 S pseudoviruses (counting ∼20,000 relative light units) were incubated with serially diluted (120-fold to 87,480-fold dilution at the final concentration) heat-inactivated sera at 37°C for 1 hour. Pseudoviruses without sera were included as controls. Then, a 40 μl mixture of pseudovirus and serum/antibody was added to HOS-ACE2/TMPRSS2 cells (10,000 cells/50 μl) in a 96-well white plate. At 2 d.p.i., the infected cells were lysed with a One-Glo luciferase assay system (Promega, Cat# E6130), a Bright-Glo luciferase assay system (Promega, Cat# E2650), or a britelite plus Reporter Gene Assay System (PerkinElmer, Cat# 6111 6066769), and the luminescent signal was measured using a GloMax explorer multimode microplate reader 3500 (Promega) or CentroXS3 (Berthhold Technologies). The assay of each serum sample was performed in triplicate, and the 50% neutralization titer (NT_50_) was calculated using Prism 9 software v9.1.1 (GraphPad Software).

### SARS-CoV-2 preparation and titration

The working virus stocks of SARS-CoV-2 were prepared and titrated as previously described^2,5,10,23–26,28,30,61^. In this study, clinical isolates of B.1.1 (strain TKYE610670; GISAID ID: EPI_ISL_479681)25, Delta (B.1.617.2, strain TKYTK1734; GISAID ID: EPI_ISL_2378732)^26^, BA.2 (strain TY40-385; GISAID ID: EPI_ISL_9595859)^5^, BA.5 (strain TKYS14631; GISAID ID: EPI_ISL_15669344) were used. In brief, 20 μl of the seed virus was inoculated into VeroE6/TMPRSS2 cells (5,000,000 cells in a T-75 flask). One h.p.i., the culture medium was replaced with DMEM (low glucose) (Wako, Cat# 041-29775) containing 2% FBS and 1% PS. At 3 d.p.i., the culture medium was harvested and centrifuged, and the supernatants were collected as the working virus stock.

The titer of the prepared working virus was measured as the 50% tissue culture infectious dose (TCID_50_). Briefly, one day before infection, VeroE6/TMPRSS2 cells (10,000 cells) were seeded into a 96-well plate. Serially diluted virus stocks were inoculated into the cells and incubated at 37°C for 4 days. The cells were observed under a microscope to judge the CPE appearance. The value of TCID_50_/ml was calculated with the Reed–Muench method^62^.

For verification of the sequences of SARS-CoV-2 working viruses, viral RNA was extracted from the working viruses using a QIAamp viral RNA mini kit (Qiagen, Cat# 52906) and viral genome sequences were analyzed as described above (see “Viral genome sequencing” section). Information on the unexpected substitutions detected is summarized in **Supplementary Table S6**, and the raw data are deposited in the Sequence Read Archive (accession ID: PRJDB14899).

### Yeast surface display

Yeast surface display (**Fig. 3a**) was performed as previously described^2, 10, 21, 22^. Briefly, the RBD genes [“construct 3” in ref.^22^, covering residues 330–528] in the pJYDC1 plasmid were cloned by restriction enzyme-free cloning and transformed into the EBY100 Saccharomyces cerevisiae. The primers are listed in **Supplementary Table S5**. The expression media 1/9 (ref.^63^) was inoculated (OD 1) by overnight (220 rpm, 30°C, SD-CAA media) grown culture, followed by cultivation for 24 hours at 20°C. The medium was supplemented with 10 mM DMSO solubilized bilirubin (Sigma-Aldrich, Cat# 14370-1G) for expression cocultivation labeling [pJYDC1, eUnaG2 reporter holo-form formation, green/yellow fluorescence (excitation at 498 nm, emission at 527 nm)]. Cells (100 µl aliquots) were collected by centrifugation (3000 g, 3 minutes), washed in ice-cold PBSB buffer (PBS with 1 mg/ml BSA), and resuspended in an analysis solution with a series of CF®640R succinimidyl ester labeled (Biotium, Cat# 92108) ACE2 peptidase domain (residues 18–740) concentrations. The peptidase domain of wild-type ACE2 and ACE2 N90Q were produced and purified as previously described^22^. The reaction volume was adjusted (1–100 ml) to avoid the ligand depletion effect, and the suspension was incubated overnight in a rotator shaker (10 rpm, 4°C). Incubated samples were washed with PBSB buffer, transferred into a 96-well plate (Thermo Fisher Scientific, Cat# 268200), and analyzed by a CytoFLEX S Flow Cytometer (Beckman Coulter, USA, Cat#. N0-V4-B2-Y4) with the gating strategy described previously^22^. The eUnaG2 signals were compensated by CytExpert software (Beckman Coulter). The mean binding signal (FL4-A) values of RBD-expressing cells, subtracted by signals of nonexpressing populations, were subjected to the determination of the dissociation constant K_D_, Y_D_ by a noncooperative Hill equation fitted by nonlinear least-squares regression using Python v3.7 fitted together with two additional parameters describing the titration curve^22^.

### Pseudovirus infection

Pseudovirus infection (**Fig. 3b**) was performed as previously described^2, 5, 10, 23, 26, 28, 31, 32, 36, 38, 58, 60^. Briefly, the amount of pseudoviruses prepared was quantified by the HiBiT assay using a Nano Glo HiBiT lytic detection system (Promega, Cat# N3040) as previously described^35, 64^. For measurement of pseudovirus infectivity, the same amount of pseudoviruses (normalized to the HiBiT value, which indicates the amount of HIV-1 p24 antigen) was inoculated into HOS-ACE2/TMPRSS2 cells, HEK293-ACE2 cells or HEK293-ACE2/TMPRSS2 cells and viral infectivity was measured as described above (see “Neutralization assay” section). For analysis of the effect of TMPRSS2 on pseudovirus infectivity (**Fig. 3c**), the fold change of the values of HEK293-ACE2/TMPRSS2 to HEK293-ACE2 was calculated.

### SARS-CoV-2 S-based fusion assay

A SARS-CoV-2 S-based fusion assay (**Fig. 3d,e**) was performed as previously described^2, 5, 10, 23–28^. Briefly, on day 1, effector cells (i.e., S-expressing cells) and target cells (Calu-3/DSP_1-7_ cells) were prepared at a density of 0.6–0.8 × 10^6^ cells in a 6-well plate. On day 2, for the preparation of effector cells, HEK293 cells were cotransfected with the S expression plasmids (400 ng) and pDSP_8-11_ (ref.^65^) (400 ng) using TransIT-LT1 (Takara, Cat# MIR2300). On day 3 (24 hours posttransfection), 16,000 effector cells were detached and reseeded into a 96-well black plate (PerkinElmer, Cat# 6005225), and target cells were reseeded at a density of 1,000,000 cells/2 ml/well in 6-well plates. On day 4 (48 hours posttransfection), target cells were incubated with EnduRen live cell substrate (Promega, Cat# E6481) for 3 hours and then detached, and 32,000 target cells were added to a 96-well plate with effector cells. *Renilla* luciferase activity was measured at the indicated time points using Centro XS3 LB960 (Berthhold Technologies). For measurement of the surface expression level of the S protein, effector cells were stained with rabbit anti-SARS-CoV-2 S S1/S2 polyclonal antibody (Thermo Fisher Scientific, Cat# PA5-112048, 1:100). Normal rabbit IgG (Southern Biotech, Cat# 0111-01, 1:100) was used as a negative control, and APC-conjugated goat anti-rabbit IgG polyclonal antibody (Jackson ImmunoResearch, Cat# 111-136-144, 1:50) was used as a secondary antibody. The surface expression level of S proteins (**Fig. 3d**) was measured using a FACS Canto II (BD Biosciences) and the data were analyzed using FlowJo software v10.7.1 (BD Biosciences). For calculation of fusion activity, *Renilla* luciferase activity was normalized to the MFI of surface S proteins. The normalized value (i.e., *Renilla* luciferase activity per the surface S MFI) is shown as fusion activity.

### AO-ALI model

An airway organoid (AO) model was generated according to our previous report^2, 10, 29, 30^. Briefly, normal human bronchial epithelial cells (NHBEs, Cat# CC-2540, Lonza) were used to generate AOs. NHBEs were suspended in 10 mg/ml cold Matrigel growth factor reduced basement membrane matrix (Corning, Cat# 354230). Fifty microliters of cell suspension were solidified on prewarmed cell culture-treated multiple dishes (24-well plates; Thermo Fisher Scientific, Cat# 142475) at 37°C for 10 min, and then, 500 μl of expansion medium was added to each well. AOs were cultured with AO expansion medium for 10 days. For maturation of the AOs, expanded AOs were cultured with AO differentiation medium for 5 days.

The AO-ALI model (**Fig. 3i**) was generated according to our previous report^10, 66^. For generation of AO-ALI, expanding AOs were dissociated into single cells, and then were seeded into Transwell inserts (Corning, Cat# 3413) in a 24-well plate. AO-ALI was cultured with AO differentiation medium for 5 days to promote their maturation. AO-ALI was infected with SARS-CoV-2 from the apical side.

### Preparation of human airway and lung epithelial cells from human iPSCs

The air-liquid interface culture of airway and lung epithelial cells (**Fig. 3j,k**) was differentiated from human iPSC-derived lung progenitor cells as previously described^5, 10, 30, 67–69^. Briefly, lung progenitor cells were induced stepwise from human iPSCs according to a 21-day and 4-step protocol^67^. At day 21, lung progenitor cells were isolated with the specific surface antigen carboxypeptidase M and seeded onto the upper chamber of a 24-well Cell Culture Insert (Falcon, #353104), followed by 28-day and 7-day differentiation of airway and lung epithelial cells, respectively. Alveolar differentiation medium with dexamethasone (Sigma-Aldrich, Cat# D4902), KGF (PeproTech, Cat# 100-19), 8-Br-cAMP (Biolog, Cat# B007), 3-isobutyl 1-methylxanthine (IBMX) (Fujifilm Wako, Cat# 095-03413), CHIR99021 (Axon Medchem, Cat# 1386), and SB431542 (Fujifilm Wako, Cat# 198-16543) was used for the induction of lung epithelial cells. PneumaCult ALI (STEMCELL Technologies, Cat# ST-05001) with heparin (Nacalai Tesque, Cat# 17513-96) and Y-27632 (LC Laboratories, Cat# Y-5301) hydrocortisone (Sigma-Aldrich, Cat# H0135) was used for induction of airway epithelial cells.

### Airway-on-a-chips

Airway-on-a-chips (**Fig. 3l,m**) were prepared as previously described^2, 10, 29, 30^. Human lung microvascular endothelial cells (HMVEC-L) were obtained from Lonza (Cat# CC-2527) and cultured with EGM-2-MV medium (Lonza, Cat# CC-3202). For preparation of the airway-on-a-chip, first, the bottom channel of a polydimethylsiloxane (PDMS) device was precoated with fibronectin (3 μg/ml, Sigma-Aldrich, Cat# F1141). The microfluidic device was generated according to our previous report^70^. HMVEC-L cells were suspended at 5,000,000 cells/ml in EGM2-MV medium. Then, 10 μl of suspension medium was injected into the fibronectin-coated bottom channel of the PDMS device. Then, the PDMS device was turned upside down and incubated. After 1 hour, the device was turned over, and the EGM2-MV medium was added into the bottom channel. After 4 days, AOs were dissociated and seeded into the top channel. AOs were generated according to our previous report^66^. AOs were dissociated into single cells and then suspended at 5,000,000 cells/ml in the AO differentiation medium. Ten microliter suspension medium was injected into the top channel. After 1 hour, the AO differentiation medium was added to the top channel. In the infection experiments (**Fig. 3l,m**), the AO differentiation medium containing either BA.2, BA.2.75, XBB or Delta isolate (500 TCID_50_) was inoculated into the top channel. At 2 h.p.i., the top and bottom channels were washed and cultured with AO differentiation and EGM2-MV medium, respectively. The culture supernatants were collected, and viral RNA was quantified using RT–qPCR (see “RT–qPCR” section above).

### Microfluidic device

A microfluidic device was generated according to our previous report^10, 70^. Briefly, the microfluidic device consisted of two layers of microchannels separated by a semipermeable membrane. The microchannel layers were fabricated from PDMS using a soft lithographic method. PDMS prepolymer (Dow Corning, Cat# SYLGARD 184) at a base to curing agent ratio of 10:1 was cast against a mold composed of SU-8 2150 (MicroChem, Cat# SU-8 2150) patterns formed on a silicon wafer. The cross-sectional size of the microchannels was 1 mm in width m in height. Access holes were punched through the PDMS using a 6-mm biopsy punch (Kai Corporation, Cat# BP-L60K) to introduce solutions into the microchannels. Two PDMS layers were bonded to a PET membrane m pores (Falcon, Cat# 353091) using a thin layer of liquid PDMS prepolymer as the mortar. PDMS prepolymer was spin-coated (4000 rpm for 60 sec) onto a glass slide. Subsequently, both the top and bottom channel layers were placed on the glass slide to transfer the thin layer of PDMS prepolymer onto the embossed PDMS surfaces. The membrane was then placed onto the bottom layer and sandwiched with the top layer. The combined layers were left at room temperature for 1 day to remove air bubbles and then placed in an oven at 60°C overnight to cure the PDMS glue. The PDMS devices were sterilized by placing them under UV light for 1 hour before the cell culture.

### SARS-CoV-2 infection

One day before infection, Vero cells (10,000 cells), VeroE6/TMPRSS2 cells (10,000 cells) and Calu-3 cells (10,000 cells) were seeded into a 96-well plate. SARS-CoV-2 [1,000 TCID50 for Vero cells (**Fig. 3f**); 100 TCID50 for VeroE6/TMPRSS2 cells (**Fig. 3g**) and Calu-3 cells (**Fig. 3h**)] was inoculated and incubated at 37°C for 1 hour. The infected cells were washed, and 180 μl of culture medium was added. The culture supernatant (10 μl) was harvested at the indicated timepoints and used for RT–qPCR to quantify the viral RNA copy number (see “RT–qPCR” section below). In the infection experiments using AO-ALI (**Fig. 3i**), human iPSC-derived airway and lung epithelial cells (**Fig. 3j,k**), working viruses were diluted with Opti-MEM (Thermo Fisher Scientific, Cat# 11058021). The diluted viruses (1,000 TCID50 in 100 μl) were inoculated onto the apical side of the culture and incubated at 37_°C for 1 _hour. The inoculated viruses were removed and washed twice with Opti-MEM. For collection of the viruses, 100 μl Opti-MEM was applied onto the apical side of the culture and incubated at 37_°C for 10_minutes. The Opti-MEM was collected and used for RT–qPCR to quantify the viral RNA copy number (see “RT–qPCR” section below). The infection experiments using an airway-on-a-chip system (**Fig. 3l,m**) were performed as described above (see “Airway-on-a-chips” section).

### RT–qPCR

RT–qPCR was performed as previously described2,5,10,23-26,28,30,61. Briefly, 5 μl culture supernatant was mixed with 5 μl of 2 × RNA lysis buffer [2% Triton X-100 (Nacalai Tesque, Cat# 35501-15), 50 mM KCl, 100 mM Tris-HCl (pH 7.4), 40% glycerol, 0.8 U/μl recombinant RNase inhibitor (Takara, Cat# 2313B)] and incubated at room temperature for 10 min. RNase-free water (90 μl) was added, and the diluted sample (2.5 μl) was used as the template for real-time RT-PCR performed according to the manufacturer’s protocol using One Step TB Green PrimeScript PLUS RT-PCR kit (Takara, Cat# RR096A) and the following primers: Forward *N*, 5’-AGC CTC TTC TCG TTC CTC ATC AC-3’; and Reverse *N*, 5’-CCG CCA TTG CCA GCC ATT C-3’. The viral RNA copy number was standardized with a SARS-CoV-2 direct detection RT-qPCR kit (Takara, Cat# RC300A). Fluorescent signals were acquired using a QuantStudio 1 Real-Time PCR system (Thermo Fisher Scientific), QuantStudio 3 Real-Time PCR system (Thermo Fisher Scientific), QuantStudio 5 Real-Time PCR system (Thermo Fisher Scientific), StepOne Plus Real-Time PCR system (Thermo Fisher Scientific), CFX Connect Real-Time PCR Detection system (Bio-Rad), Eco Real-Time PCR System (Illumina), qTOWER3 G Real-Time System (Analytik Jena), Thermal Cycler Dice Real Time System III (Takara) or 7500 Real-Time PCR System (Thermo Fisher Scientific).

### Animal experiments

Animal experiments (**Fig. 4 and Extended Data Fig. 3**) were performed as previously described^2, 5, 10, 23, 25, 26, 30^. Syrian hamsters (male, 4 weeks old) were purchased from Japan SLC Inc. (Shizuoka, Japan). For the virus infection experiments, hamsters were anesthetized by intramuscular injection of a mixture of 0.15 mg/kg medetomidine hydrochloride (Domitor^®^, Nippon Zenyaku Kogyo), 2.0 mg/kg midazolam (Dormicum^®^, Fujifilm Wako, Cat# 135-13791) and 2.5 mg/kg butorphanol (Vetorphale^®^, Meiji Seika Pharma) or 0.15 mg/kg medetomidine hydrochloride, 4.0 mg/kg alphaxaone (Alfaxan^®^, Jurox) and 2.5 mg/kg butorphanol. Delta, BA.2.75 and XBB (10,000 TCID_50_ in 100 µl) or saline (100 µl) was intranasally inoculated under anesthesia. Oral swabs were collected at the indicated timepoints. Body weight was recorded daily by 7 d.p.i. Enhanced pause (Penh), the ratio of time to peak expiratory follow relative to the total expiratory time (Rpef) were measured every day until 7 d.p.i. (see below). Lung tissues were anatomically collected at 2 and 5 d.p.i. The viral RNA load in the oral swabs and respiratory tissues was determined by RT–qPCR. These tissues were also used for IHC and histopathological analyses (see below).

### Lung function test

Lung function tests (**Fig. 4a**) were routinely performed as previously described^2, 5, 10, 23, 25, 26^. The two respiratory parameters (Penh and Rpef) were measured by using a Buxco Small Animal Whole Body Plethysmography system (DSI) according to the manufacturer’s instructions. In brief, a hamster was placed in an unrestrained plethysmography chamber and allowed to acclimatize for 30 seconds. Then, data were acquired over a 2.5-minute period by using FinePointe Station and Review software v2.9.2.12849 (DSI).

### Immunohistochemistry

Immunohistochemistry (IHC) (**Fig. 4c and Extended Data Fig. 3**) was performed as previously described^2, 5, 10, 23, 25, 26^ using an Autostainer Link 48 (Dako). The deparaffinized sections were exposed to EnVision FLEX target retrieval solution high pH (Agilent, Cat# K8004) for 20 minutes at 97°C for activation, and a mouse anti-SARS-CoV-2 N monoclonal antibody (clone 1035111, R&D Systems, Cat# MAB10474-SP, 1:400) was used as a primary antibody. The sections were sensitized using EnVision FLEX for 15 minutes and visualized by peroxidase-based enzymatic reaction with 3,3’-diaminobenzidine tetrahydrochloride (Dako, Cat# DM827) as substrate for 5 minutes. The N protein positivity was evaluated by certificated pathologists as previously described^2, 5, 10, 23, 25, 26^. Images were incorporated as virtual slides by NDP.scan software v3.2.4 (Hamamatsu Photonics). The N-protein positivity was measured as the area using Fiji software v2.2.0 (ImageJ).

### H&E staining

H&E staining (**Fig. 4d**) was performed as previously described^2, 5, 10, 23, 25, 26^. Briefly, excised animal tissues were fixed with 10% formalin neutral buffer solution and processed for paraffin embedding. The paraffin blocks were sectioned at a thickness of 3 µm and then mounted on MAS-GP-coated glass slides (Matsunami Glass, Cat# S9901). H&E staining was performed according to a standard protocol.

### Histopathological scoring

Histopathological scoring (**Fig. 4e**) was performed as previously described^2, 5, 10, 23, 25, 26^. The inflammation area in the infected lungs was measured by the presence of the type II pneumocyte hyperplasia. Four hamsters infected with each virus were sacrificed on days 2 and 5 d.p.i., and all four lung lobes, including right upper (anterior/cranial), middle, lower (posterior/caudal), and accessory lobes, were sectioned along with their bronchi. The tissue sections were stained by H&E, and the digital microscopic images were incorporated into virtual slides using NDRscan3.2 software (Hamamatsu Photonics). The color of the images was decomposed by RGB in split channels using Fiji software v2.2.0.

Histopathological scoring was performed as described in the previous studies^2, 5, 10, 23, 25, 26^. Pathological features including bronchitis or bronchiolitis, hemorrhage or congestion, alveolar damage with epithelial apoptosis and macrophage infiltration, hyperplasia of type II pneumocytes, and the area of the hyperplasia of large type II pneumocytes were evaluated by certified pathologists and the degree of these pathological findings were arbitrarily scored using four-tiered system as 0 (negative), 1 (weak), 2 (moderate), and 3 (severe). The “large type II pneumocytes” are the hyperplasia of type II pneumocytes μm-diameter nucleus. Total histology score is the sum of these five indices. In the representative lobe of each lung, the inflammation area with type II pneumocytes was gated by the certificated pathologists on H&E staining, and the indicated area were measured by Fiji software v2.2.0.

### Statistics and reproducibility

Statistical significance was tested using a two-sided Mann–Whitney *U* test, a two-sided Student’s *t* test, a two-sided Welch’s *t* test, or a two-sided paired *t-*test unless otherwise noted. The tests above were performed using Prism 9 software v9.1.1 (GraphPad Software).

In the time-course experiments (**Figures 3e–l, 4a–4c, and 4e, and S2b**), a multiple regression analysis including experimental conditions (i.e., the types of infected viruses) as explanatory variables and timepoints as qualitative control variables was performed to evaluate the difference between experimental conditions thorough all timepoints. The initial time point was removed from the analysis. The *P* value was calculated by a two-sided Wald test. Subsequently, familywise error rates (FWERs) were calculated by the Holm method. These analyses were performed in R v4.1.2 (https://www.r-project.org/).

Principal component analysis to representing the antigenicity of the S proteins was performed (**Fig. 2d**). The NT50 values for biological replicates were scaled, and subsequently, principal component analysis was performed using the prcomp function on R v4.1.2 (https://www.r-project.org/).

In **Figures 5D, 5F and S5**, photographs shown are the representative areas of at least two independent experiments by using four hamsters at each timepoint. In **Figure S4D**, photographs shown are the representatives of >20 fields of view taken for each sample.

## Data availability

All databases/datasets used in this study are available from the GISAID database (https://www.gisaid.org) and GenBank database (https://www.gisaid.org; EPI_SET ID: EPI_SET_221223pb, EPI_SET_221223ew, EPI_SET_221223yk, EPI_SET_221222mt). Viral genome sequencing data for working viral stocks are available in the Sequence Read Archive (accession ID:

## Code availability

The computational codes used in the present study and the GISAID supplemental tables for EPI_SET_221223pb, EPI_SET_221223ew, EPI_SET_221223yk, EPI_SET_221222mt are available in the GitHub repository (https://github.com/TheSatoLab/XBB).

