## Supplementary figures and images for "Virological characteristics of the SARS-CoV-2 XBB variant derived from recombination of two Omicron subvariants"

### Figure S1

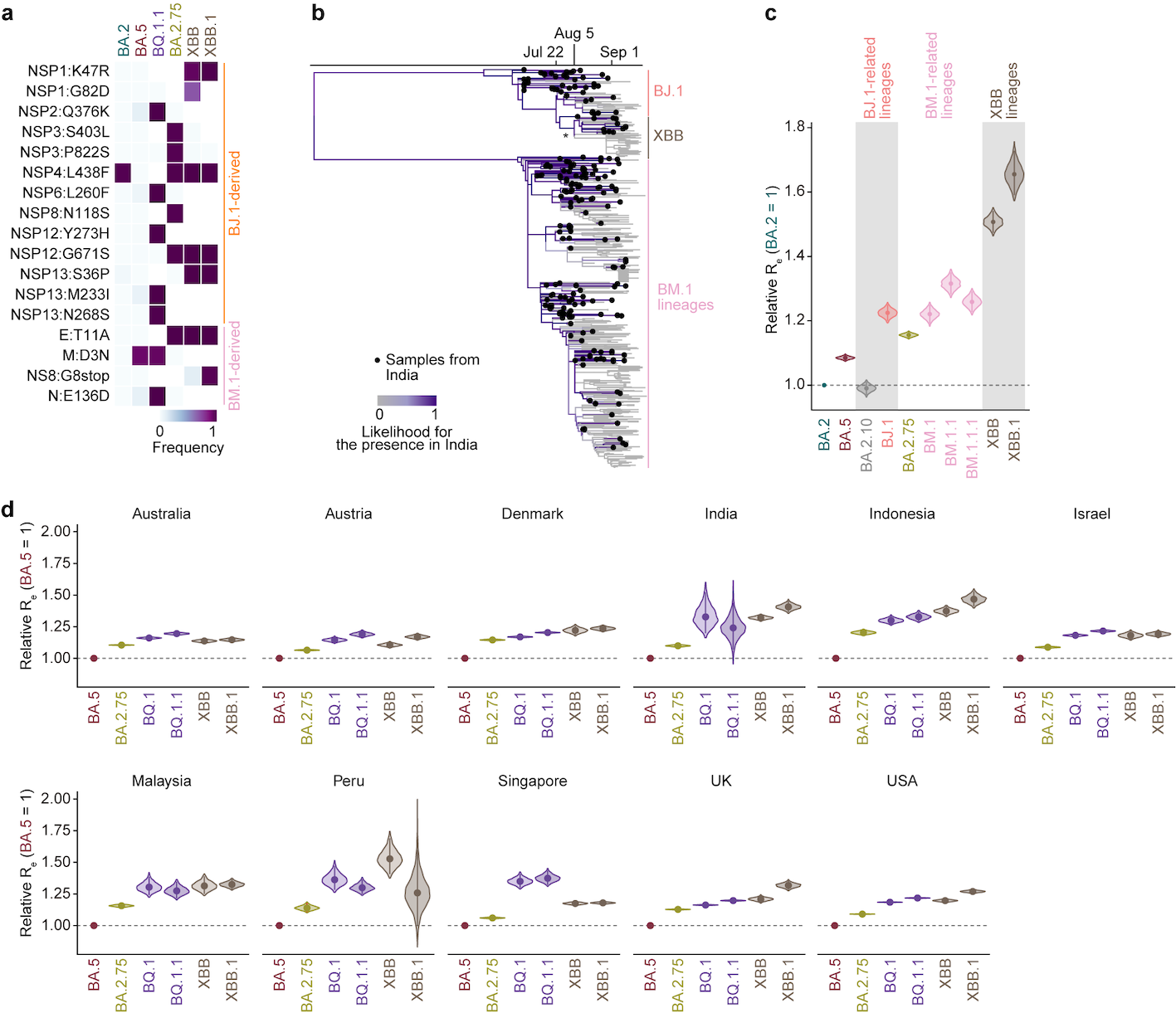

### Figure S2

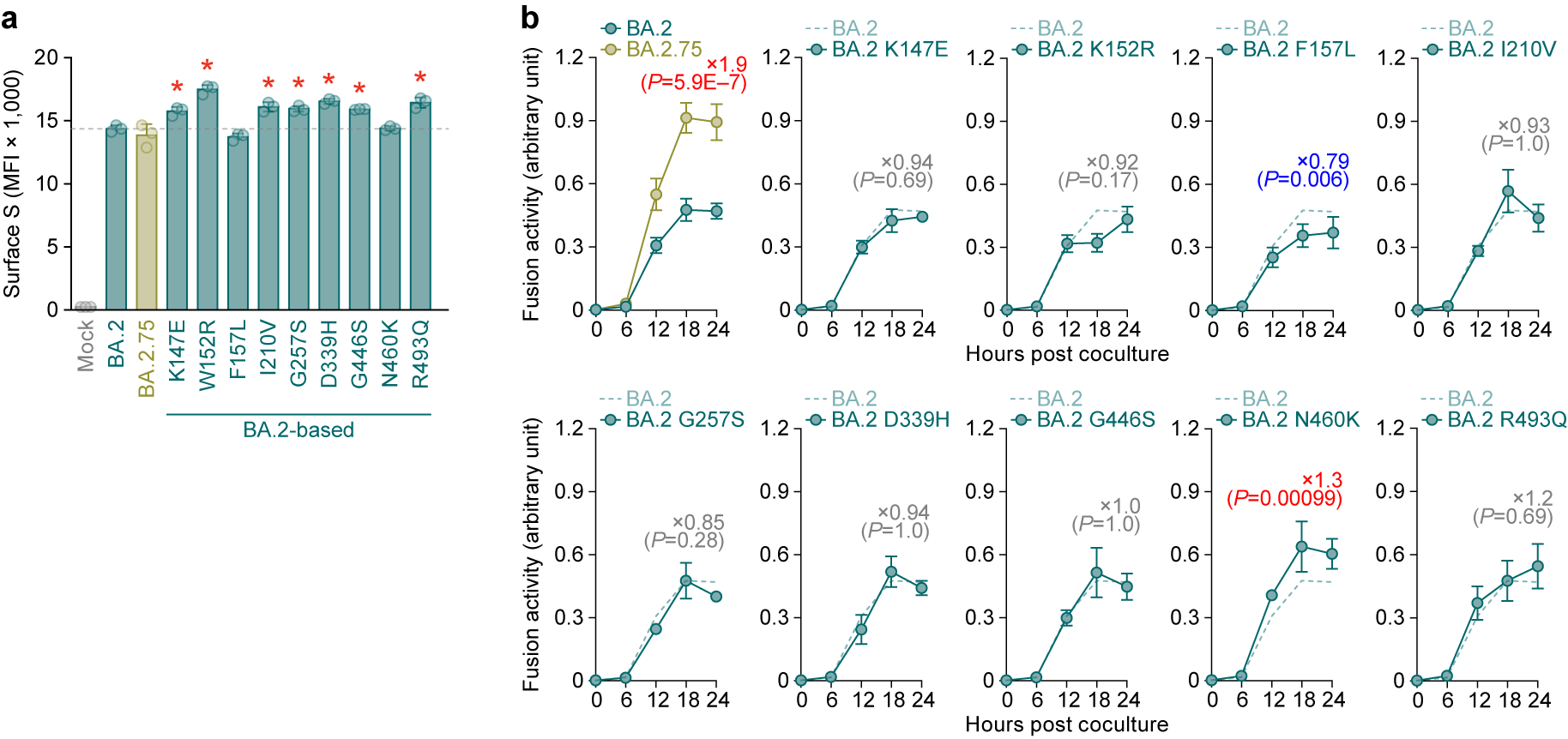

### Figure S3

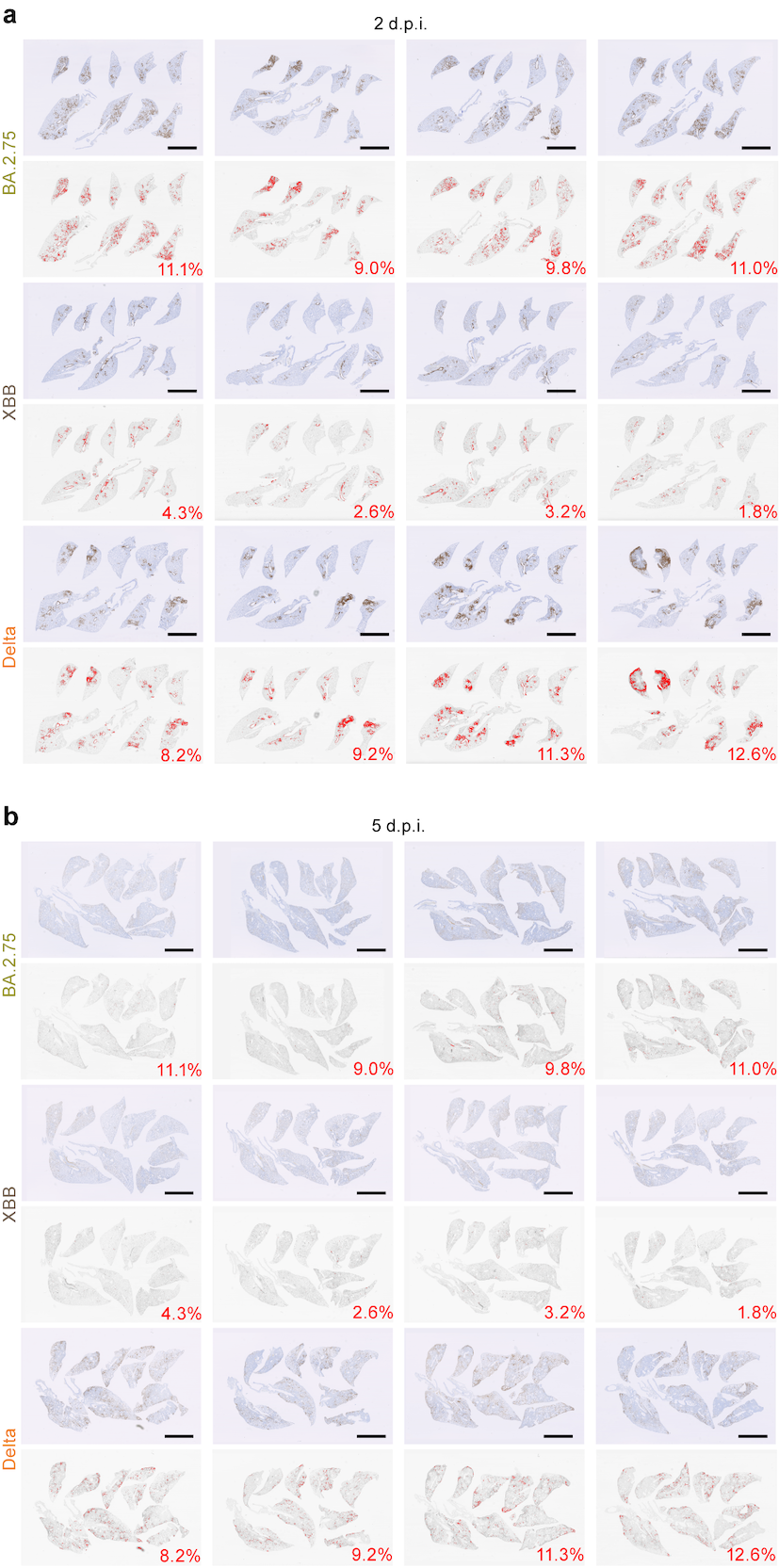
